# Defining the spatial distribution of extracellular adenosine revealed a myeloid-dependent immunosuppressive microenvironment in pancreatic ductal adenocarcinoma

**DOI:** 10.1101/2022.05.24.493238

**Authors:** Vincenzo Graziano, Andreas Dannhorn, Heather Hulme, Kate Williamson, Hannah Buckley, Saadia A Karim, Sheng Y. Lee, Brajesh P. Kaistha, Sabita Islam, James E. D. Thaventhiran, Frances M. Richards, Richard Goodwin, Rebecca Brais, Jennifer P Morton, Simon J. Dovedi, Alwin G. Schuller, Jim Eyles, Duncan I. Jodrell

## Abstract

The prognosis for patients with pancreatic ductal adenocarcinoma (PDAC) remains extremely poor. It has been suggested that the adenosine pathway contributes to the ability of PDAC to evade the immune system and its resistance to immunotherapies (Immuno-Oncology Therapy, IOT), by generating extracellular adenosine (eAdo).

Using genetically engineered allograft models of PDAC in syngeneic mice with differential immune infiltration and response to IOT, we showed enrichment of the adenosine pathway in tumour-infiltrating immune cells (in particular, myeloid populations). Using MS-Imaging, we showed that extracellular adenosine distribution is heterogeneous in tumours, with high concentrations in hypoxic margins that surround necrotic areas, associated with a rich myeloid infiltration, demonstrated using Imaging Mass Cytometry (IMC). Pro-tumorigenic M2 macrophages express high levels of the Adora2a receptor; particularly in the IOT resistant model. Blocking the *in vivo* formation and function of eAdo (Adoi), using a combination of anti-CD73 antibody and an Adora2a inhibitor slowed tumour growth and reduced metastatic burden. In addition, blocking the adenosine pathway improved the efficacy of combinations of cytotoxic agents or immunotherapy. Finally, Adoi remodelled the tumour microenvironment (TME), as evidenced by reduced infiltration of M2 macrophages and Tregs. RNAseq analysis showed that genes related to immune modulation, hypoxia and tumour stroma were downregulated following Adoi and a specific adenosine signature derived from this is associated with a poorer prognosis in patients with PDAC.

The formation of eAdo appears to promote the development of the immunosuppressive TME in PDAC, contributing to its resistance to conventional and novel therapies. Therefore, inhibition of the adenosine pathway may represent a strategy to modulate the stroma and improve therapy response in patients with PDAC.

## Introduction

Survival for patients with pancreatic ductal adenocarcinoma (PDAC) has not changed significantly in the last 50 years and remains poor (https://www.cancerresearchuk.org/health-professional/cancer-statistics-for-the-uk). There is a need for new treatments, given that current standard of care for patients with metastatic disease is associated with poor outcomes, with less than 10% of patients living for more than 2 years ^1^. In addition to relative resistance to conventional therapies, cancer immunotherapy (Immuno-Oncology Therapy, IOT) is also ineffective in this disease, except in a small group of patients (1-2%) with microsatellite instability/mismatch repair deficient (MSI-H/dMMR) tumours ^2^. Several authors consider that the reason for this resistance can be ascribed to the low mutational burden of this neoplasm, which leads to lymphocyte exclusion and anergy ^3, 4^. However, the tumour microenvironment in PDAC has also been shown to be populated by a rich variety of immune cells, most of which demonstrate immune suppressive features, which contribute to the resistance to immunotherapy ^5^.

The adenosine pathway is an immunosuppressive axis which has gained much attention in cancer immunology for its role in suppressing the immune activation associated with cytotoxic treatments (chemotherapy, targeted therapy and radiotherapy) ^6^. This has led to the clinical evaluation of inhibitors of the pathway in combination with more conventional approaches ^7^. The adenosine pathway involves conversion of extracellular ATP (eATP), a powerful immune activator, to extracellular adenosine (eAdo) by the ectonucleotidases CD39 and CD73 ^8^. eADO has been linked to cancer in several studies that have demonstrated that its concentration in different tumour tissues is several folds higher than in normal tissues ^6, 9^. CD39 is overexpressed in a subpopulation of exhausted tumour-infiltrating T-cells ^10, 11^ and its expression correlates with another marker of immunosuppression (PD1 expression) ^11^. CD39 and CD73 have roles in the aggressiveness of adult glioblastoma ^12^, where they are expressed on infiltrating macrophages ^13, 14^. The adenosine signature recently published by Sidders and colleagues ^15^ shows that this pathway correlates with resistance to immunotherapies and is associated with other genetic features of tumour aggressiveness, such as p53 mutations. The abundant presence of eAdo in the microenvironment can dampen immune activation through the stimulation of a pro-tumorigenic stroma. This is mostly orchestrated by macrophages and myeloid derived suppressive cells (MDSCs) ^16, 17^, favouring a tolerogenic function of dendritic cells (DCs) ^18, 19, 20^ which results in inhibition of T-cells/NK cell activation ^21, 22^.

The myeloid populations play a pivotal role in the aggressiveness of many cancer types and in particular, PDAC. For instance, the presence of pro-tumorigenic populations of macrophages ^23, 24^ and myeloid-derived suppressive cells (MDSCs) ^25^ infiltrating the microenvironment, is associated with poor survival and correlates with immune exclusion of PDAC. Macrophages can elicit the secretion of cytokines which can on the one hand, favour the proliferation and invasiveness of cancer cells while interacting with cancer-associated fibroblasts ^26, 27, 28^, and on the other hand induce anergy and physical exclusion of adaptive immune cells ^29, 30, 31^. Targeting macrophages in a pre-clinical pancreatic cancer model has been demonstrated to be effective to obtain tumour regression and reduce metastatic formation ^32^. Unfortunately, this approach has not translated into clinical benefit, which in part can be explained by the fact that the global reduction of the tumour-infiltrating macrophages can be biologically different from reprogramming distinct tumour associated macrophage (TAM) subtypes ^28^.

Some recent publications link the adenosine pathway to the biology and aggressiveness of PDAC. PDAC has been shown to have an increased adenosine pathway RNA signature associated with a worse prognosis ^15^, and genes encoding for the receptors for eAdo as well as CD73 have been found to be overexpressed in bulk-RNA sequencing (RNAseq) when comparing tumours to normal pancreatic tissue ^33^. However, little is known about the complex mechanism generated by the adenosine pathway resulting in the immunosuppressive characteristics of pancreatic cancer microenvironment and stroma, in particular the role that the adenosine pathway has in shaping the immune infiltration of this disease.

Here, we propose a model where the tumour-infiltrating immune cell populations of PDAC generate an axis of immunosuppression, where extracellular adenosine produced mostly in hypoxic regions of the tumour (identified using Mass Spectrometry Imaging), enriched for the myeloid cell infiltration, stimulates pro-tumorigenic M2 macrophages. The axis described is expressed preferentially by the IOT-resistant model when compared to the IOT-responsive one. Therefore, blocking the adenosine pathway in the IOT-resistant PDAC model, strongly suppresses the formation of extracellular adenosine and reshapes the immune microenvironment, favouring disease control when combined with cytotoxic treatments and immunotherapies. Bulk RNAseq gene analysis confirms the role of the myeloid-dependent adenosine pathway in PDAC survival, underpinning the importance of our results for understanding the biological complexity and the clinical utility of the adenosine pathway inhibition.

## Methods and materials

### Cell lines and chemicals

*Kras*^LSL-G12D/+^; *Trp53*^LSL-R172H/+^; *Pdx1*-Cre; *Rosa26*^YFP/YFP^ (KPCY)-derived cell lines 2838c3, 6499c4, 6620c1 (IOT-responsive), 6419c5, 6694c2 and 6422c1 (IOT-resistant) were a kind gift from Ben Stanger (University of Pennsylvania). The cell lines were obtained from single cell cloning strategy, as described previously, and were generated from tumours developed in KPCY mice on a C57BL/6 background ^34^. Cells were grown up to 20 passages in DMEM (with pyruvate, L-glutamine and D-glucose; Gibco, #41966029) supplemented with 5% FBS (Gibco, #10270106). All the cell lines were analysed for STR fingerprinting and tested for mycoplasma routinely.

### Mice and *in vivo* experiments

Tumour allograft experiments were performed in the animal facility (Biological Resource Unit, BRU) of the CRUK Cambridge Institute, in accordance with the UK Animals Scientific Procedures Act 1986, with approval from the CRUK Cambridge Institute Animal Ethical Review and Welfare Body. 8-12 week old female C57BL/6 mice were used for *in vivo* experiments and were purchased from Charles River (UK).

Tumour allograft studies were performed with technical assistance from CRUK-CI BRU staff. Mice were subcutaneously injected in the right flank with 1×10^6^ KPCY-derived cells in 50% PBS and 50% Matrigel basement membrane matrix (#354234, Corning). In the interventional experiments, mice were treated as indicated, starting 12-14 days from tumour cell implantation, to allow the microenvironment to establish. Tumour volume was calculated using the formula; (π/6)*(width)^2^*length. Tumour response was defined based on the % of change of the longest diameter from start of therapy (stable disease < 20% increase and < 30% decrease of target lesion RECIST v. 1.1). Mice were then killed at specific endpoints (e.g. 14 days from start of treatment) or when the tumour reached 2000 mm^3^ (or before in case of appearance of clinical signs).

*Kras*^LSL-G12D/+^; *Trp53*^LSL-R172H/+^; *Pdx1*-Cre (KPC) mice for tumour phenotyping, were obtained from a breeding colony maintained by the CRUK-CI Genome Editing Core team. Tumours were detected by palpation followed by ultrasound imaging by the Genome Editing Core. Tumour tissues, spleens and mesenteric and inguinal lymph nodes from KPC mice were provided once tumour dimensions or health status rendered them unsuitable for therapeutic studies. KPC mice were killed when showing clinical signs of the disease (swollen abdomen, loss of body conditioning resembling cachexia, reduced mobility).

KPC mice used for interventional study were bred at the CRUK Beatson Institute and maintained on a mixed background. All work was performed under UK Home Office license and approved by the University of Glasgow Animal Welfare and Ethical Review Board. Mice of both sexes, in similar proportions, were used in all cohorts. Mice suspected to have PDAC following palpation were anesthetised in 0.2L/min medical air and isoflurane and then underwent 3D ultrasound imaging using the VisualSonics Vevo 3100 ultrasound system with MX550D 40μm resolution transducer (FujiFilm). Mice were randomly assigned to 1 of the 4 treatment groups (A: vehicles + isotype; B: AZD6738 + gemcitabine; C: AZD4635 + 2c5mIgG1; D: AZD6738 + gemcitabine + AZD4635 + 2c5mIgG1) once tumours were confirmed by imaging, and follow-up scans were performed weekly until endpoint was reached. Schedule of the treatments are specified in the results and figures sections. Mice were culled when exhibiting moderate symptoms of pancreatic cancer (see above). Statistical assessment of survival was carried out by Kaplan-Meier and Log-Rank analysis. Analysis of ultrasound images was performed using VevoLab software (version 3.1.1) from VisualSonics. In 3D mode, stacked images of the tumour were imported and the tumour border annotated, allowing a 3D construct to be formed.

When indicated the following drugs were used: AZD6738 (ATRi; 25 mg/kg daily for 4 days), AZD4635 (Adora2ai; 50 mg/kg bid), 2c5mIgG1 (anti-CD73; 10 mg/kg twice weekly), AB740080 D265A (anti-PD-L1: 10 mg/kg twice weekly), NIP228 mouse IgG1 control kappa (isotype; 10 mg/kg twice weekly) and NIP228 muIgG1 D265A (isotype; 10 mg/kg twice weekly) were provided by AstraZeneca; gemcitabine hydrochloride (Tocris, 3259) was used at 100 mg/kg twice weekly; inVivoPlus anti-CD40 (clone FGK4.5/FGK45; bioxcell BE0016-2) and InVivoPlus rat IgG2a isotype control, anti-trinitrophenol (clone 2A3; bioxcell BE0089) were used as a single injection of 100 μg. InVivoPlus anti-CTLA-4 (clone 9H10; bioxcell BP0131) or InVivoPlus isotype control polyclonal Syrian hamster IgG (bioxcell BP0087): 200 μg/dose × 3 times.

### Clonogenic assay

KPCY-derived cells were seeded at 200 cells/well in a 6-well plate. After 24 hours, cells were treated with different concentrations of anti-CD73 (2c5mIgG1) antibody (1, 10, 100 μg/ml) or isotype (NIP228 mouse IgG1 control kappa) for 8 days. Antibodies were added every 3 days. On day 8, colonies were stained with SRB protocol previously described (36). Images were taken and analysed using GelCount (Oxford Optronix). Colony forming efficiency was calculated as a ratio between the number of colonies and number of plated cells. Surviving fractions were calculated as the ratio between wells treated with anti-CD73 antibody and the ones treated with isotype. At least 3 wells per condition were plated for each of the 3 replicates per experiment.

### IncuCyte time lapse imaging

KPCY-derived cells were plated at 2500 cells/well density in a 96-well black-wall plate (at least 3 wells per condition). Cells were grown in cell culture medium supplemented with the indicated concentration of anti-CD73 (2c5mIgG1) or isotype (NIP228 mouse IgG1 control kappa) antibody in triplicate. Images were acquired with 10x objective, every 3 hours from 3 different fields per well using Incucyte Live cells imaging microscope (Essen Bioscience). Confluence was calculated as the average of the 3 fields using the Incucyte algorithm. Experiments were repeated at least 3 times.

### Single cell suspension preparation

For experiments in subcutaneous allografts, tumours were weighed and placed in RPMI and finely minced with scissors in a 2 ml tube, which was then washed with up to 2.5 ml of digestion buffer (Tumour dissociation kit, Miltenyi, 130-096-730) plus Deoxyribonuclease I (300 μg/ml, Sigma, DN25-1G). Dissociation was performed using the protocol suggested by Miltenyi. For KPC tumours, a trypsin inhibitor (250 μg/ml, Sigma, T6522) was added to the digestion buffer. Following the digestion, the samples were passed through a 70 μM strainer filter (Grainer Bio-one, 542-070), washing with MACS buffer (PBS + 0.1% FBS and 2nM EDTA).

Mouse spleens, inguinal and mesenteric lymph nodes were mashed on a 100 μM filter (Grainer Bio-one, 542-000) over a 50 ml tube, using a syringe plug and the filter was washed with MACS buffer and centrifuged (at 4° C as for all the following centrifugation). Red cell lysis buffer (1 ml; 0.15M Ammonium Chloride; 10mM Potassium hydrogen carbonate; 0.1mM EDTA; pH 7.4 (adjusted with KOH) was then used to resuspend splenocytes, 3 minutes at room temperature and then washed with MACS buffer and centrifuged at 300g for 5 minutes. Samples were eventually resuspended in 200-400 μl of MACS buffer.

### Flow cytometry

Single cell suspensions were aliquoted in a round-bottom 96 well/plate (Costar, 3879) and stained with live/dead fixable stain (Invitrogen, L34962; 1:100 in PBS) for 10 minutes at room temperature. After washing in MACS buffer, cells were FC-blocked with anti-CD16/32 antibody (BioLegend Cat# 101320, RRID:AB_1574975; 1:100) for 5 minutes. Then, antibodies for surface staining were added and incubated at 4° C degrees for 30 minutes. After washing, cells were fixed with FACS fix buffer [PBS + 1% Formaldehyde + 0.02 g/ml Glucose + 0.02% Sodium Azide] for 10 minutes and then washed and resuspended in MACS buffer for FACS analysis. For intracellular staining, cells were fixed with fixation/permeabilization buffer (Invitrogen 00-5123-43 and 00-5223-56) for 15 minutes, then washed in perm buffer and stained with the relevant antibody in perm buffer for 60 minutes. Cells were then washed and resuspended in MACS buffer for FACS analysis. Samples were acquired using BD-Symphony flow cytometer and the generated FCS files were analysed using FlowJo V10 software (RRID:SCR_008520). The following antibodies were used and gating strategies are shown in the supplementary figures(myeloid populations and lymphoid population; suppl. fig. 1A and 1B respectively): BV786-CD45 (BD Biosciences Cat# 564225, RRID:AB_2716861; 1:200), APC/Fire750-CD3 (BioLegend Cat# 100248, RRID:AB_2572118; 1:50), BV650-CD8 (BioLegend Cat# 100741, RRID:AB_11124344; 1:100), BV711-CD4 (BioLegend Cat# 100549, RRID:AB_11219396; 1:200), APC-Foxp3 (Invitrogen Cat# 17-5773-82, RRID:AB_469457; 1:100), FITC-CD19 (BioLegend Cat# 115505, RRID:AB_313640; 1:200), BV510-CD11b (BioLegend Cat# 101245, RRID:AB_2561390; 1:200), PerCP/Cy5.5-CD44 (BioLegend Cat# 103031, RRID:AB_2076206; 1:200), BV421-PD1 (BioLegend Cat# 135218, RRID:AB_2561447; 1:100), BV421-PD-L1 (BioLegend Cat# 124315, RRID:AB_10897097; 1:100), PE/Cy7-CD39 (BioLegend Cat# 143805, RRID:AB_2563393; 1:100), PE-CD73 (BioLegend Cat# 127206, RRID:AB_2154094; 1:100), FITC-F4/80 (BioLegend Cat# 123108, RRID:AB_893502; 1:200), PE/Cy7-CD206 (BioLegend Cat# 141719, RRID:AB_2562247; 1:100), PerCP/Cy5.5-Ly6C (BioLegend Cat# 128012, RRID:AB_1659241; 1:100), APC/Cy7-Ly6G (BioLegend Cat# 127624, RRID:AB_10640819; 1:100), AF700-MHCII (BioLegend Cat# 107629, RRID:AB_2290801; 1:200), PE-CD11c (eBioscience Cat# 12-0114-83, RRID:AB_465553; 1:200), BV605-NKp46 (BioLegend Cat# 137619, RRID:AB_2562452; 1:25), APC-Adora2a (Novus Biotech Cat# NBP1-39474APC; 1:150), APC-CD86 (BioLegend Cat# 105012, RRID:AB_493342; 1:100). Gating strategy for immune subpopulations is shown in supplementary figure 1A-B. For CD73 *in vitro* staining of KPCY-derived cell lines, cells were stained as described above with live/dead fixable stain and PE-CD73 antibody and analysed with BD-Symphony flow cytometer. For in vitro treatment, cells were treated with anti-CD73 (2c5mIgG1) or isotype (NIP228) antibody at a concentration of 10 μg/ml for 24 hours. Competitive staining was performed before this experiment to confirm there was no competition between 2c5mIgG1 and TY11.8 (PE-CD73) clones (data not shown).

### Tissue preparation for Mass Spectrometry Imaging (MSI) and Imaging Mass Cytometry (IMC) analysis

PDAC mouse tumours were snap frozen in liquid nitrogen immediately after resection and the tissues were embedded in a HPMC/PVP hydrogel as previously described ^35^. Sectioning was performed on a CM3050 S cryostat (Leica Biosystems, Nussloch, Germany) at a section thickness of 10 μm and the tissue sections were immediately thaw mounted and dried under a stream of nitrogen and sealed in vacuum pouches to preserve the metabolic integrity of the sections. Tissue sections for DESI-MSI and IMC were thaw-mounted onto Superfrost microscope slides (Thermo Scientific Waltham, MA, USA), whilst sections prepared for MALDI-MSI were thaw mounted onto conductive ITO coated slides (Bruker Daltonik, Bremen, Germany). Polyvinylpyrrolidone (PVP) (MW 360 kDa) and (Hydroxypropyl)-methylcellulose (HPMC) (viscosity 40-60 cP, 2 % in H2O (20 C) were purchased from Merck (Darmstadt, Germany). Methanol, water, iso-pentane and isopropyl alcohol were obtained from Fisher Scientific (Waltham, MA, USA).

### Mass Spectrometry Imaging (MSI)

DESI-MSI analysis was performed on a Q-Exactive mass spectrometer (Thermo Scientific, Bremen, Germany) equipped with an automated 2D-DESI ion source (Prosolia Inc., Indianapolis, IN, USA) operated in negative ion mode, covering the applicable mass range up to a m/z of 1000, with a nominal mass resolution of 70,000. The injection time was fixed to 150 ms resulting in a scan rate of 3.8 pixel/s. The spatial resolution was adapted between experiments to allow acquisition of the data for all directly compared samples within a single experiment of 48 h, with pixel sizes ranging from 65-75 μm. A home-built Swagelok DESI sprayer was operated with a mixture of 95% methanol, 5% water delivered with a flow rate of 1.5 μL/min and nebulized with nitrogen at a backpressure of 6 bar. The resulting .raw files were converted into .mzML files using ProteoWizard msConvert ^36^ (version 3.0.4043) and subsequently compiled to an .imzML file (imzML converter ^37^ version 1.3). All subsequent data processing was performed in SCiLS Lab (version 2021b, Bruker Daltonik, Bremen, Germany).

MALDI-MSI analysis was performed on a RapifleX Tissuetyper instrument (Bruker Daltonik, Bremen, Germany) operated in negative detection mode. 9-Aminoacridine (9-AA) prepared in 80:20 methanol:water was used as a MALDI matrix and spray deposited using an automated spray system (M3-Sprayer, HTX technologies, Chapel Hill, NC, USA). MALDI experiments were performed with a spatial resolution of 50 μm. A total of 400 laser shots were summed up per pixel to give the final spectra. For all experiments the laser was operated with a repetition rate of 10 kHz. All raw data were directly uploaded and processed in SCiLS lab (Version 2021b) software packages. All DESI and MALDI data and images were normalised to the total ion current (TIC) to compensate for signal variation across the course of the experiments. Data segmentation pipeline is shown in supplementary materials and methods.

### Imaging Mass Cytometry (IMC)

Imaging mass cytometry was performed on a slide which had already been analysed by DESI-MSI. Antibodies used for IMC staining are shown in Table 1. Untagged antibodies were tagged in house, using Fluidigm Maxpar Antibody Labelling Kit, according to manufacturer’s instructions. Following DESI-MSI analysis, the slide was fixed with 4% paraformaldehyde in phosphate-buffered saline (PBS) for 10 minutes. The slide was washed 3 × 5 minutes in PBS, permeabilized for 5 minutes with 1:1000 dilution of Triton X-100 in Casein Solution, washed 3 × 5 minutes in PBS, and blocked for 30 minutes with Casein Solution. Antibodies were diluted to an appropriate concentration in Casein Solution and the slide incubated overnight with the antibody solution at 4 °C. The slide was washed 3 × 5 minutes in PBS and nuclei were stained with DNA Intercalator-iridium at a dilution of 1:400 in PBS for 30 minutes. The slide was washed 3 × 5 minutes in PBS, 30 seconds in deionized water, then dried for storage at room temperature until analysis. A region for IMC analysis was selected using consecutive H and E stained sections and the DESI-MSI results. A box with approximately 2 × 1.8 mm area was selected for analysis to include necrotic, necrotic margin and viable tumour regions. IMC analysis was performed using a Hyperion instrument (Fluidigm Corporation, San Francisco, CA, USA) with an ablation energy of 6 db an ablation frequency of 200 Hz. IMC images were produced using MCD viewer (Version 1.0, Fluidigm) and analysis was performed using HALO (Indica labs). Tissue regions were classified using random forest with all markers included. Cells positive for each marker were manually optimised by setting a cell intensity threshold. Values for the numbers of positive cells for markers of interest were exported for analysis in Graphpad Prism v.8 (RRID:SCR_002798)

### Immunohistochemistry (IHC)

Immunohistochemistry was performed as previously described^38^ in the histopathology core at the CRUK CI. Briefly, tissues were removed from the mouse at the endpoint and immediately formalin-fixed for 24 hours. Fixed tissues were then processed, embedded in paraffin and sectioned (3 μm sections). Following dewaxing and rehydration, as standard, antigen retrieval was performed using Leica’s Epitope Retrieval Solution 2 (Tris EDTA) at 100° C for 20 minutes. Additional protein block from Dako (X090930-2) was applied. The staining using anti-mouse CD8 (Cell signalling, #98941), anti-mouse Foxp3 (Affymetrix. #14-5773) and anti-mouse p53 (Novocastra; #NCL-L-p53-CM5p) antibodies, was performed on Leica’s automated Bond-III platform in conjunction with their Polymer Refine Detection System (DS9800) and a modified version of their standard template. Slides were dehydrated and cleared in xylene on Leica’s automated ST5020 before sections were mounted on Leica’s coverslipper, CV5030 (mounting media: DPX Mountant for Histology; Sigma Aldrich, 06522-500ML) and scanned using a ScanScopeAT2 (Aperio Leica Biosystems). Quantification of viable tumour tissue was performed after exclusion of necrotic area using the Halo software v. 3.3.2541.405 (Indica Labs). Cell density was calculated as the number of positive cells x mm2 of tumour tissue analysed. Sections of mouse spleen were used on each slide as internal control.

For the analysis of the lung metastatic burden of any individual mouse, the 4 right lobes and the left lobe were cut into multiple pieces and together fixed and then embedded, then treated as above. p53 staining was used for helping the detection of smaller lesions (min. of 5 cells). Analysis was performed using Halo software and expressed as % of metastatic areas/total lung area analysed. Mice with intra-abdominal/thoracic organs direct infiltration were excluded from the analysis.

### RNA sequencing

RNA was extracted from twelve subcutaneous allograft tumour tissues (6 mice of the vehicle + isotype group or control and 6 mice of the AZD4635 + 2c5mIgG1 group or Adoi) weighing up to 30 mg. Tissues were firstly disrupted and homogenised using TissueLyser II and then RNA was extracted using a Qiagen RNAeasy kit, according to the manufacturer’s instructions. RNA was then quantified using a Qubit 3.0 (life technologies) and purity and quality were assessed using an Agilent 4150 (G2992AA) TapeStation system (Agilent). Library construction was followed by paired-end 50 bp sequencing on a Novaseq 6000 sequencer.

### Bioinformatics analysis

Sequencing files in FASTQ format were aligned against the GRCh38 mouse genome using HISAT2 (RRID:SCR_015530) with default parameters. Samtools (RRID:SCR_002105) was used to create, index and merge BAM files of reads from different lanes belonging to individual samples. FeatureCounts (RRID:SCR_012919) was utilized to quantify gene-level expression of transcripts. All downstream analyses were completed in R version 4.1.2. Prior to analysis, MSI data for sequenced samples were examined. From the vehicle/isotype-treated arm (control group), sample 23729 showed minimal necrosis, low peri-necrotic adenosine and a high ATP/AMP ratio suggesting a very high energetic state. This identified the sample as a potential outlier which was confirmed upon visual inspection of a PCA plot (Supplementary fig. 1C). It was excluded prior to downstream analysis.

For the remaining 11 samples, initially genes were filtered to maintain only genes that were expressed at a reasonable level in >5 treatment conditions using the filterByExpr() command from the edgeR package (ver 3.36.0).

Differential gene expression analyses were performed on raw read counts of the combined data object of all 11 samples. To identify significantly expressed genes between Adoi and control groups, we utilised a Wald test within the DESeq2 package (RRID:SCR_000154, ver 1.34.0). Genes were considered differentially expressed when the analysis resulted in an adjusted P-value (corrected for multiple testing using the Benjamini and Hochberg method) below 0.05. The volcano plot was generated using the EnhancedVolcano package (ver 1.12.0) with the addition of custom code.

Gene set enrichment analysis (GSEA) of 712 genes identified as differentially expressed with padj≤0.05 and log2(fold change)≤-0.58 was performed via the Enrichr (RRID:SCR_001575) (suppl. table 2) server database for Kyoto Encyclopedia of Genes and Genomes (KEGG) (https://www.genome.jp/kegg/) pathways and Gene Ontology (GO): Biological Processes (http://geneontology.org/). Subsequently, enriched terms ranked for significance for each database were downloaded and are reported in Supplementary Table 4-5. Terms of interest were selected from the top 15 ranks in each table. Genes from this study which were shown to be enriched in these terms of interest were then selected to be displayed in a heatmap. Raw counts were normalised with DESeq2 (RRID:SCR_000154) prior to visualisation of gene expression levels with pheatmap (ver 1.0.12). Please refer to the supplementary methods and materials references section for all of the above.

### Analysis of human PDAC available datasets and generation of PDAC-specific adenosine signature

In order to the evaluate the correlation of the adenosine-related gene expression profile to human PDAC we analysed 712 genes which had at least a 50% decreased expression (Log2FC <-0.58) following adenosine inhibition treatment, of which 561 had a human ortholog (suppl. table 2-3).

For the analysis of the adenosine-related gene expression in Bailey ^39^ PDAC subtypes (ADEX, Immunogenic, Squamous and Pancreatic progenitor) we derived z-scores for the 517 genes analysed in the dataset (out of the 561 genes) for the 97 patients with RNAseq data and subtype information (https://www.cbioportal.org/, RRID:SCR_014555). The z-score of all genes were summed per patient and the total number represented as the adenosine pathway gene score as previously shown ^15^ and in supplementary materials and methods.

For the generation of a PDAC-specific adenosine signature and application of this to PDAC survival, from the list of 561 human ortholog genes, we manually curated the ones related without ambiguity to the major biological processes implicated in PDAC pathogenesis and indicated by pathway analysis (hypoxia, immunity and extracellular matrix organisation). Of these genes, only those that correlated positively or negatively to survival in PDAC (https://kmplot.com) and were significantly co-expressed with CD73 and/or Adora2a in public datasets (Bailey et al. or TCGA) were selected. A final list of 52 genes was analysed (suppl. table 6).

Using PDAC specific data from TCGA ^40^ available in https://www.cbioportal.org/, we derived the z-score of these 52 genes for each patient with known disease-specific-survival (DSS), Progression free survival (PFS) and disease-free-survival (DFS). The z-scores for all genes were summed up for each patient and was deemed high adenosine score if >0 or low adenosine score if <0, as previously shown ^15^.

### Statistics

Graphpad Prism v.8 (RRID:SCR_002798) was used for statistical analyses. Analysis and comparisons of two groups was performed with two-tailed unpaired Student’s T-test when assuming Gaussian distribution or Mann Whitney test. Analysis of three or more groups was performed with one-way ANOVA with Tukey’s multiple comparisons post-test analysis unless otherwise specified. Kaplan-Meier analysis with log-rank Mantel-Cox test was used to evaluate difference in survivals. Differences were considered significant when p<0.05.

### Data Availability Statement

The data generated in this study are available within the article and its supplementary data files or from the corresponding authors upon reasonable request. Code for differential expression analysis and visualisation of RNAseq data is available via Github https://github.com/ka-lw/AdenoPDAC.

## Results

### Adenosine pathway expression on KPCY-derived cell lines

Human PDAC cells have been shown to express for CD73 expression and demonstrate sensitivity, although weakly, to the targeting of CD73 *in vitro* ^41^. For this reason, we sought to investigate whether murine KPCY-derived cell lines (which are associated with contrasting ability to generate IOT-resistant or responsive tumours when re-implanted in syngeneic mice) express the proteins of the canonical adenosine pathway (CD39, CD73 and Adora2a). We found that, as in human cells, mouse PDAC cell lines express CD73 (from 72% to 99% of cells; fig. 1A and suppl. fig. 2A) but demonstrate negligible or no expression of CD39 and Adora2a (suppl. fig. 2B-C). Exposing cells to an anti-CD73 antibody (2c5mIgG1) reduced significantly the detection of CD73 in all the cells after only 24-hour treatment (p<0.05 in all cell lines, fig. 1B), but this did not translate into inhibition of cell growth after short or long exposure at high concentrations. In addition, confluency experiments showed that the treatment did not affect the proliferation of any of the cell lines over a period of 72 hours (fig. 1C and suppl. fig. 2D-E), and colony-forming experiments performed on 2838c3 (IOT-responsive) and 6419c5 (IOT-resistant) demonstrated no differences in terms of number or size of the colonies formed after 8 days of continuous treatment (fig. 1D-E and suppl. fig. 2F). In order to evaluate whether a direct effect of anti-CD73 exposure affects cell proliferation, reducing adenosine formation, we cultured KPCY-derived cell lines with increasing concentrations of AMP and 5’-N-(Ethylcarboxamido)adenosine (NECA, a stable form of adenosine). We were able to demonstrate that adenosine and AMP have no effect on the proliferation capacity of these cell lines (suppl. fig. 2G-H), corroborating the hypothesis of a non-cancer cell direct effect of anti-CD73 therapeutic approaches.

**Figure 1.**
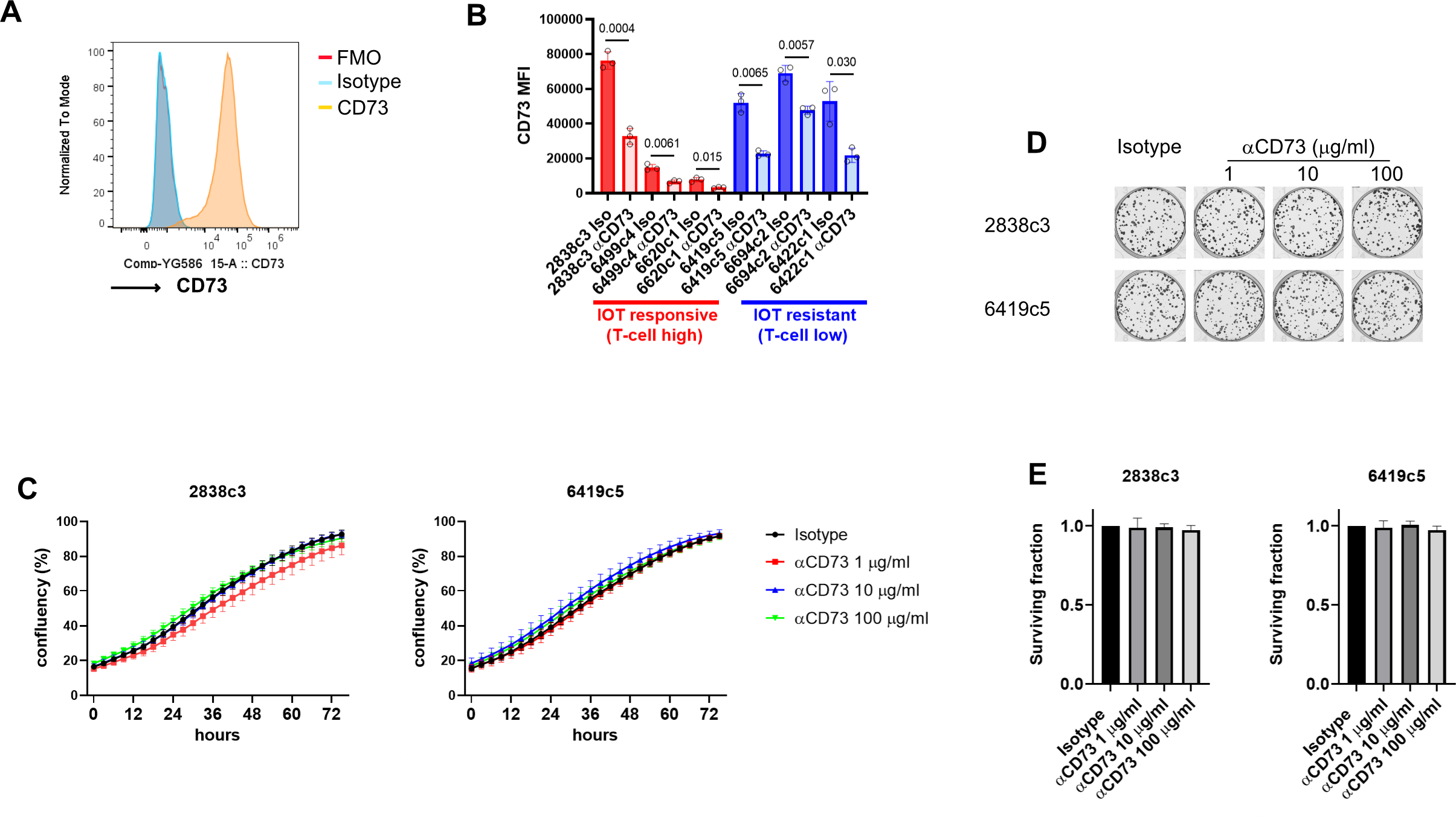
Expression of CD73 on KPCY-derived cell lines and response to anti-CD73 in vitro inhibition. (A) Representative histogram of CD73 expression on KPCY-derived cell line (2838c3) in flow cytometry. (B) CD73 expression was evaluated on KPCY-derived cell lines after 24 hours incubation with 10 μg/ml of anti-CD73 or isotype. (C) 2838c3 (left) and 6419c5 (right) cell lines were grown with increasing concentration of anti-CD73 or isotype (100 μg/ml) and confluency was evaluated using IncuCyte time lapse imaging for up to 72 hours. For each experiment, 3 different wells per condition were used per experiment. (D-E) Representative images (D) and graphs (E) showing survival fraction of cells (2838c3 left, 6419c5 right) from the colony-forming experiment following 8-day treatment with anti-CD73 or isotype. For each experiment, 3 different wells per condition were used. All data are presented as mean ± SEM from experiments repeated 3 times. Statistical analysis was performed with two-tailed unpaired Student’s t-test (B), mixed-effect model (C) and one-way ANOVA with post-test analysis for multiple comparisons; p values are shown in the graphs when considered significant (p<0.05).

### Comparison between KPCY-derived cell lines allograft and KPC autochthonous tumours

Despite the lack of activity in our cell line experiments, preliminary reports from clinical trials suggest activity in patients with PDAC treated with an anti-CD73 antibody (oleclumab) ^42^. As we hypothesised that this might be the result of impact on the tumour microenvironment (TME), we investigated the expression of this pathway on the tumour-infiltrating immune cells for different murine PDAC models, including KPCY-derived cell line allografts (with differential response to IOT) and autochthonous KPC tumours. In order to understand the complexity and similarities of the immune system in these models we first compared the immune infiltration of the cell line allografts to the KPC model.

The immunosuppressive characteristics identified in autochthonous KPC tumours, appear to be more aligned with those of the IOT-resistant model. In particular regarding lymphocyte populations, KPC tumours are usually infiltrated by a low number of CD8^+^ T cells (mean number of CD45^+^ cells for 2838c3 is 6.9%, 0.9% for 6419c5 and 1.9% for KPC tumours) which bear fewer features of activation/exhaustion (mean 70% vs 13% vs 10%) as shown in suppl. figure 3A and have a similar total CD4^+^ T-cells (suppl. fig. 3C) and regulatory T-cell (Tregs) infiltration (mean 3,2% vs 1% vs 0.7%, suppl. fig 3B) to IOT-resistant tumours. Moreover, KPC tumours showed greater heterogeneity regarding myeloid infiltrating populations (suppl. fig 3D-H). These results suggest that our IOT-resistant and responsive models stand out as the extreme clonotypes which can arise from the complex and heterogeneous biology found in KPC autochthonous tumours.

### The adenosine pathway is enriched in immune cells infiltrating PDAC models

We hypothesized that the adenosine pathway might have a more impactful role in the TME, as opposed to a cell autonomous effect. For this reason, we investigated the expression of the adenosine pathway components on tumour-infiltrating immune cells which represent significant proportion cells seen in PDAC lesions. We showed a highly significant enrichment in both the IOT-resistant and IOT-responsive models for CD39^+^CD73^+^ double-expressing immune cells, when compared to secondary lymphoid organs (spleen and nodes). In particular, the majority (65-91% in tumour vs 36-56% in spleen) of tumour-infiltrating CD11b^+^ myeloid cells express the two receptors, due to an increase in expression of CD73, given that those cells are normally CD39^+^ (fig. 2A-B). Similar results were shown for Tregs and CD8^+^ T-cells, which are normally CD73^+^ and displayed an increase in expression of CD39 in tumour, compared to the secondary lymphoid organs counterparts (suppl. fig. 4A; p<0.05). There was no significant difference in these findings when comparing the two models, despite their differential response to IOT. We then confirmed these findings in KPC autochthonous tumours and found similar results, with a significant increase of CD39-CD73 double expressing CD11b^+^ myeloid cells infiltrating the tumours compared to spleens (mean 77% vs 36% p<0.0001, fig. 2C-D), harvested from the same mice. Of note, 4 KPC mice had synchronous metastases (3 liver and 1 spleen), and in 3 of these, myeloid cells infiltrating the metastases were also enriched for CD73^+^CD39^+^ double expression (suppl. fig. 4 C) when compared to secondary lymphoid organs. Significant increased co-expression was also observed for Tregs and CD8^+^ T-cells in KPC tumours when compared to mesenteric or inguinal lymph nodes (suppl. fig. 4B). An increased percentage of CD39/CD73 double expressing CD8 T-cells and Tregs was noted in the spleens (suppl. fig. 4B) if compared to what was found in non-tumour bearing mice (not shown) or subcutaneous tumour bearing mice, suggesting trafficking of immune lymphoid populations between primary lesions and closer lymphoid organs.

**Figure 2.**
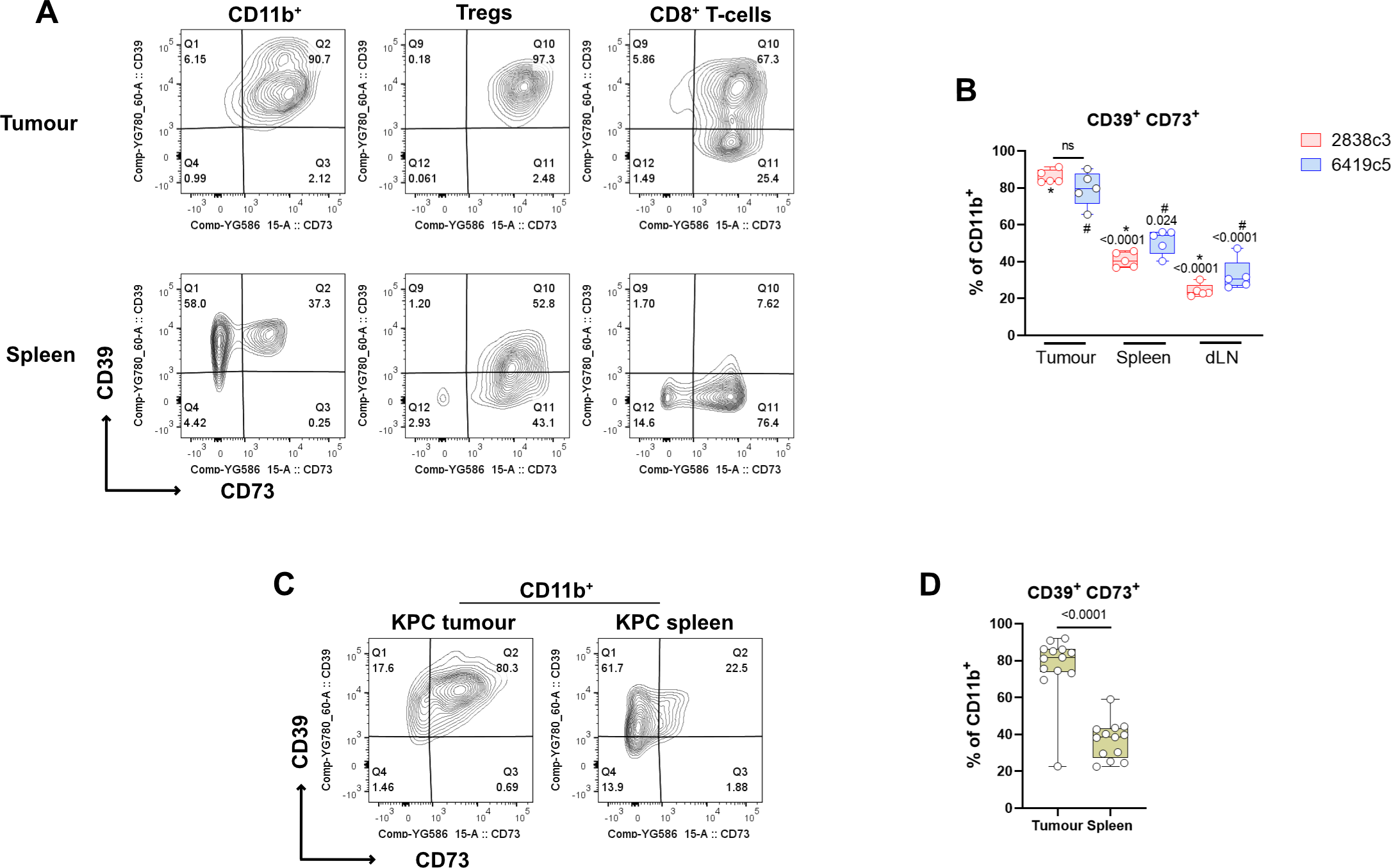
The adenosine pathway members are expressed on PDAC-infiltrating immune cells. (A) Representative flow cytometry plots showing expression of CD39 and CD73 on myeloid population (left), Tregs (middle) and CD8+ T-cells (right) from KPCY-cell line derived tumour (upper) and matched spleen (lower) (N= 5 mice per group). (B) Box and whisker graph showing CD39^+^CD73^+^ double expression on CD11b^+^ cells for 2838c3 (N=5) and 6419c5 (N=5) in tumours, matched spleens and tumour draining lymph nodes. (C-D) Representative flow cytometry plots (C) and box and whisker graph (D) showing CD39^+^CD73^+^ double expression on CD11b^+^ cells infiltrating autochthonous KPC tumours (N=13). All data are presented as interleaved box and whiskers. Statistical analysis was performed using one-way ANOVA with post-hoc test analysis for multiple comparisons (B) and two-tailed unpaired Student’s t-test (D); p values are shown in the graphs when considered significant (p<0.05).

### Adenosine distributes primarily in the areas surrounding necrosis

Given the enrichment of the adenosine pathway in the TME of PDAC models, we anticipated that extracellular adenosine might have been abundant in the tumour microenvironment. Using Mass Spectrometry Imaging (MSI) we evaluated the presence and the distribution of the purinergic system in the TME of IOT-resistant tumours. The tissue classification and segmentation approaches were driven by tissue-defining metabolic pattern. Areas characterised by a high energetic state defined by a high abundance of ATP and ADP and a low abundance of depleted high energy phosphates such as AMP, were called *viable tumour*. In contrast, areas of tumour adjacent to necrosis (termed *necrotic margin)* were characterised by high abundance of lactate, products of ribonucleotide catabolism (i.e. xanthine and hypoxanthine) and other metabolites associated with tissue hypoxia and an overall energy-deprived state (fig. 3A-B). We found that adenosine is present in high concentrations in the microenvironment of IOT-resistant tumours, although showing a heterogenous distribution, with high abundance in the necrotic margin areas (fig. 3A,C). When we investigated the cell population distribution in the different areas using Imaging Mass Cytometry (IMC) in IOT-resistant tumours, we noted in the necrotic margin areas a 2.7-fold increase in the number of infiltrating CD11b^+^ myeloid cells (mean 1970 in the *necrotic margin* vs 730 in *viable tumour* CD11b^+^ cells), that led to a significant decrease of the ratio between cancer cells/myeloid cells (fig. 3D). This again suggests that myeloid cells have an instrumental role in the generation of adenosine in this aggressive model of PDAC.

**Figure 3.**
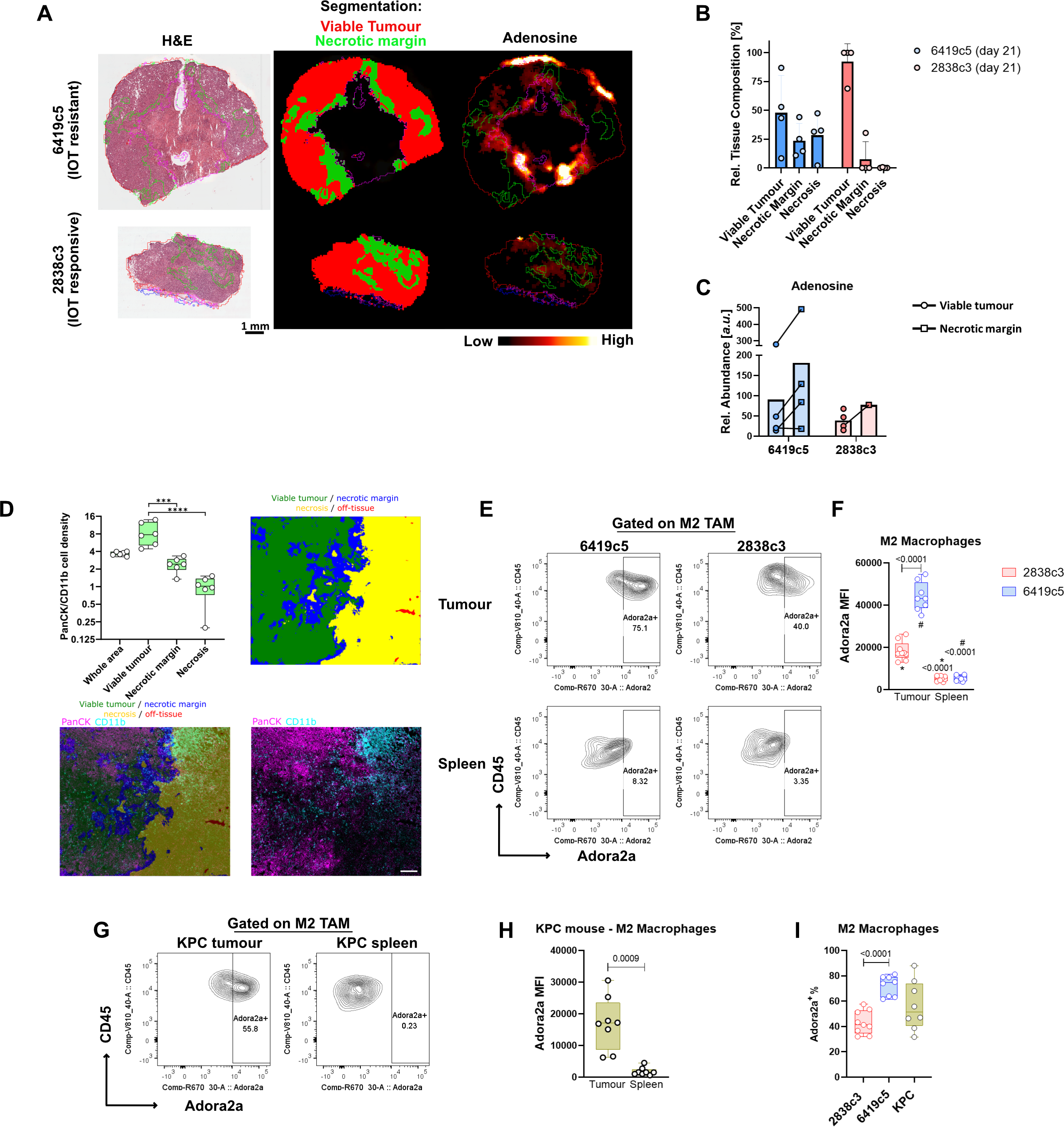
Adenosine distribution is spatially heterogeneous and targets myeloid subpopulations. (A) Mass spectrometry Imaging (MSI) representative images showing adenosine expression and distribution in PDAC allografts (4 mice per group) at day 21 post-implantation. Classification was obtained based on metabolites expression and is represented as follows: viable tumour (red), necrotic margins (green). (B) Relative tissue composition differences of viable tumour, necrotic margin and necrotic areas for 6419c5 (N=4) and 2838c3 (N=4) allografts at day 21 post-implantation. As shown, necrosis was present in only one 2838c3 sample at 21 days post-implantation. Error bars represent Standard Deviation. (C) MSI analysis showing relative abundance (a.u.) of adenosine in the different areas in 6419c5 and 2838c3 allografts at day 21 post-implantation. Bars represent means. (D) The plot (upper left) shows cell density of the ratio cancer cells (PanCK^+^, purple) / myeloid cells (CD11b^+^, blue) on a Log2 scaled plot from Imaging Mass cytometry (IMC). The results of the comparison between the whole area analysed and the other tissue regions are not shown. N = 6. Asterisks ****, show adjusted p value of <0.0001. The IMC and tissue segmentation images (upper right and lowers) are from one representative 6419c5 allograft replicate at day 28 post-implantation. The tissue segmentation of the IMC image was performed by Random Forest Classification using all markers analysed. Scale bar on the IMC image is 200 μm. Segmentation shows viable tumour (green), necrosis (yellow), necrotic margin (blue) and off-tissue (red). (E-F) Representative plots (E) and summary graph (F) from flow cytometry analysis showing Adora2a expression on pro-tumorigenic M2 macrophages in allografts (upper) derived from 6419c5 (left) and 2838c3 (right) implantation. Same expression is shown in M2 macrophages in matched spleens (lower) (8-9 mice per group were used). (G-H) Flow cytometry plots (G) and graph (H) showing expression of Adora2a in M2 macrophages comparing KPC (n=8) autochthonous tumours and matched spleens. (I) Box and whisker plot of the percentage of M2 macrophages positive for Adora2a comparing two allografts (6419c5 and 2838c3) and KPC tumours. Statistical analysis was performed using one-way ANOVA with post-hoc test analysis for multiple comparisons (D,F,I) and two-tailed unpaired Student’s t-test (H); p values are shown in the graphs when considered significant (p<0.05).

### Expression of Adora2a receptor on myeloid subpopulations of pancreatic cancer models

Having shown in these models that in the PDAC microenvironment, immunosuppressive adenosine is present abundantly, we then investigated which cells within the microenvironment might be responsive to this. We investigated the expression of the adenosine A2a receptor (Adora2a, the receptor with the highest affinity for adenosine) that has been found to be frequently overexpressed in human tumours, by flow cytometry We found that Adora2a was highly expressed by tumour-infiltrating myeloid population when compared to the spleen (suppl. fig. 4D-E) and this expression was significantly higher in the IOT-resistant model in terms of MFI (10000 vs 6700, p<0.0001) and % of Adora2a^+^ myeloid cells (15% vs 11%, p=0.009). When comparing different subpopulations, pro-tumorigenic M2 macrophages, infiltrating both IOT-resistant and IOT-responsive PDAC showed high positivity for the receptor. The IOT-resistant model had higher expression of Adora2a compared to the IOT-responsive model (fig. 3E-F; p<0.0001) and percentage of Adora2a^+^ M2 positivity [72% vs 43%; p<0.0001) (fig. 3I)]. Once more, these findings were confirmed in KPC tumours where Adora2a was found to be increased in M2 macrophages infiltrating the lesions when compared to matched spleens (fig. 3G-H). The KPC model demonstrated once again the heterogeneity of pancreatic lesions, which in terms of M2 macrophages, positive for the Adora2a receptor, covers the entire range of expression seen in the two subcutaneous models used (fig. 3I). Notably, of three KPC mice where metastatic nodules were found, Adora2a expression was found retained in the M2 macrophages infiltrating the secondary lesions (suppl. fig. 4F).

In addition to pro-tumorigenic macrophages, Adora2a expression was found enriched in other myeloid immune populations infiltrating the tumours. In particular CD11b^−^ dendritic cells, CD11b^+^ dendritic cells (suppl. fig. 4G), M1 macrophages (suppl. fig. 4H), and mo-MDSCs (suppl. fig. 4I) for both models and gMDSCs for IOT-resistant tumours (suppl. fig. 4J) express significantly higher Adora2a amount when compared to matched spleens. This expression differs significantly between IOT-responsive and resistant models in CD11b^+^ dendritic cells (mean MFI 2080 ± 174 vs 3190 ± 636 respectively; p=0.007), M1 macrophages (4520 ± 983 vs 6940 ± 1690; p=0.02) and mo-MDSCs (1890 ± 479 vs 6110 ± 1870; p=0.001) (suppl. fig. 4G-I).

### Targeting adenosine pathway delays tumour growth of murine pancreatic cancer representing a combinational therapeutic opportunity

Our data suggest a mechanism by which the myeloid population contributes to the pro-tumorigenic functionality of the pancreatic cancer microenvironment, where eAdo generated by the myeloid cell populations and cancer cells would target and stimulate further the myeloid cell subpopulations, in particular the pro-tumorigenic M2 macrophages. Therefore, we inhibited *in vivo* eAdo formation and function, using an antibody against CD73 (2c5mIgG1) and a small molecule inhibitor of Adora2a (AZD4635), a combination (Adoi) which would maximise the inhibition of the axis. The 14-day treatment was started after the microenvironment was allowed to establish (12-14 days after implantation) in the IOT-resistant allografts (fig 4A). The anti-CD73 was extremely effective in reducing the expression of CD73 on the surface of all live cells (suppl. fig. 5A). MSI data confirmed marked reduction in adenosine formation in the TME (Fig. 4 B-C). In particular, adenosine was completely abolished in the viable tumour areas, while a small amount remained in the necrotic margins, accounting for a 95% decrease (fig. 4C), highlighting the importance to inhibit not only the formation of adenosine through CD73 inhibition, but also targeting adenosine receptors. The effectiveness of the treatment on the extracellular purinergic pathway was also supported by the decrease of molecules downstream of adenosine (adenine and inosine in viable tumour and necrotic margin areas) (suppl. fig 5B-C), and the increase of upstream and alternative pathway molecules as AMP (in the necrotic margin, suppl. fig 5D) and xanthine (in both viable tumour and necrotic margin, suppl. fig 5E) respectively. There was no change in the distribution of ATP, ADP and hypoxanthine (suppl. fig 5F-H).

**Figure 4.**
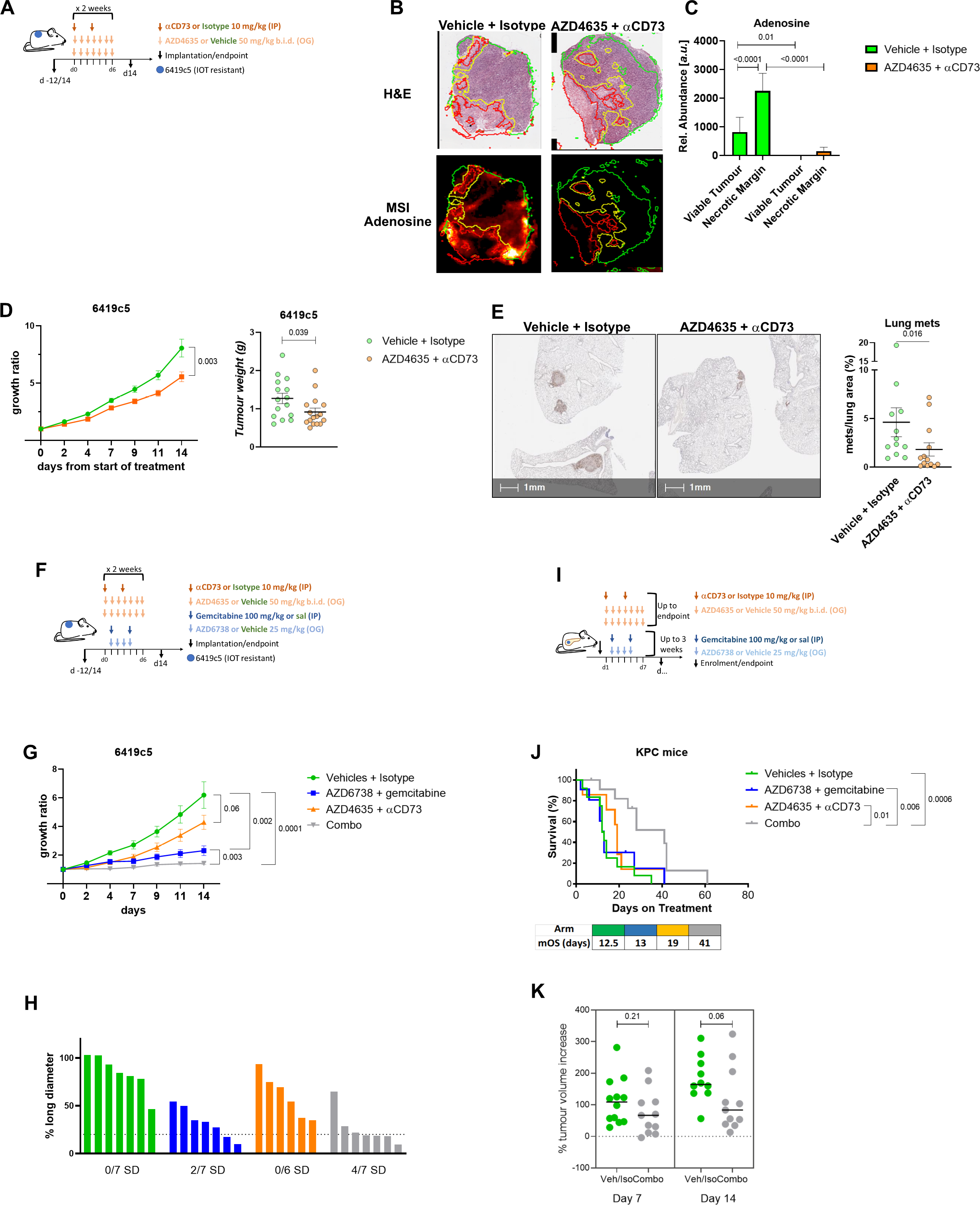
*In vivo* modulation of the adenosine pathway reduces tumour growth and metastasis and improves the efficacy of cytotoxic treatment. (A) Schedule of adenosine inhibition (Adoi) treatment. Treatment was started following 12-14 days from implantation and continued for 2 weeks. Antibody anti-CD73 (2c5mIgG1, murine IgG1) was dosed twice per week intraperitoneally at 10 mg/kg. Adora2a inhibitor (AZD4635) was given by oral gavage twice daily at 50 mg/kg. (B-C) Mass spectrometry Imaging (MSI) representative images (B) and MSI analysis graph with relative abundance (a.u.) (C) showing adenosine expression and distribution in PDAC allografts treated with vehicle + isotype or Adoi, at day 14 from treatment start (6 mice per treatment group). Classification was obtained based on metabolites expression and is represented as follows: viable tumour (green line), necrotic margins (yellow line). (D) Tumour growth ratio (left) and weight (right) of 6419c5 allografts in C57Bl/6 mice treated with anti-CD73 + AZD4635 (N=16) or vehicle + isotype (N=15). (E) Representative image (left) and graph (right) showing % of area of the lung analysed occupied by metastasis (met/lung areas x 100) in vehicle + isotype (N=12) and anti-CD73+AZD4635 (N=13), evaluated for the presence of spontaneous occurrence of lung metastases. Every dot represents a single mouse. Tissues were stained with an anti-p53 antibody to highlight the presence of cancer cells. A group of more than 5 p53-positive cells was counted as metastasis. (F) Schedule of 6419c5 tumour allografts 14-day treatment as following (N=7 mice per group): vehicles + isotype, AZD6738 + gemcitabine, anti-CD73 + AZD4635, AZD6738 + gemcitabine + anti-CD73 + AZD4635. (G) Tumour growth ratio (14 days) of 6419c5 tumour allografts treated as above (H) Percentage change in the long diameter length following 14 days of treatment per group. Number of mice with stable disease (SD, <20% increase and <30% decrease) are shown at the bottom. (I) Schedule of KPC mice treatment as following: vehicles + isotype (12 mice), AZD6738 + gemcitabine (11 mice), anti-CD73 + AZD4635 (7 mice), AZD6738 + gemcitabine + anti-CD73 + AZD4635 (combo, 12 mice). Adenosine inhibition was administered until endpoint, while AZD6738 + gemcitabine was allowed up to 3 weeks. (J) Survival analysis of KPC mice treated as above, including median overall survival (mOS). (K) Percentage of tumour volume increased in vehicles vs combo group at day 7 and 14 of treatment. All data are presented as mean ± SEM. Statistical analysis was performed using Mann-Whitney test (D-E,K), mixed-effect model (F) and one-way ANOVA with post-hoc test analysis for multiple comparisons (C). Log-rank Mantel-Cox test to evaluate difference in survivals (J); p values are shown in the graphs and considered significant when p<0.05.

The Adoi approach led to a 30% reduction of tumour growth ratio (mean of 8 vs 5.5 -folds increase from the baseline, p=0.003, fig. 4D) and tumour weight (median 1.24 vs 0.78 grams, 0.039 fig. 4A). Single agent AZD4635 induced a similar tumour control to the full combination. However, the therapeutic effect was delayed (suppl. fig. 6B), with a 30% growth reduction in the combination arm when compared to Adora2ai inhibition alone with single agent AZD4635, over the first 4 days of treatment (p=0.01). For biological reasons, given also that AZD4635 is a competitive inhibitor and its efficacy is dependent on the amount of extracellular adenosine, we chose to use it in combination with anti-CD73 antibody to maximise the blocking on extracellular adenosine effects. These data support previous findings showing the same combination had greater anti-tumour immune effect ^43^. The adenosine pathway has been shown to control the metastatic process and several authors have shown that inhibiting this axis can reduce the metastatic burden in mouse models ^44, 45^. However, we were able to show for the first time that blocking adenosine generation and function can significantly reduce the occurrence of spontaneous metastasis. The 6419c5 s.c. model spontaneously develops lung metastases in 100% of the mice and blocking adenosine strongly reduced the metastatic burden (median of % mets/lung area 0.77% vs 2.6%; p=0.016; fig. 4E).

These data suggest that targeting myeloid related, extracellular adenosine formation and effect would have an effect on tumour growth, making this approach a candidate for combinatorial therapeutic studies. Indeed, when combined with cytotoxic treatment (AZD6738 and gemcitabine; fig. 4F-G) or IOT (anti-CD40 agonist, anti-PD-L1 and anti-CTLA4 (FCP); suppl. fig. 6C), the adenosine modulation reduced further the tumour growth rate of the aggressive IOT-resistant 6419c5 tumour model. AZD6738/gemcitabine alone was able to significantly slow the growth of the IOT-resistant model (2/7 stable disease, SD [28.5%]), but the addition of the adenosine blocking (AZD4635 + 2c5mIgG1) led to further stabilization of the tumour growth in a 2-week regimen (p=0.003 vs AZD6738/gem alone; 4/7 SD [57.1%]; fig. 4G-H). These data supported the investigation of the same combination in autochthonous tumours in KPC mice (fig. 4I) to assess whether the addition of Adoi to cytotoxic treatment (AZD6738+gemcitabine) prolonged survival in this model, considered a gold standard in this disease. As figure 4J shows, the combination of Adoi and ATRi/gem induced a 3-fold increase in median overall survival (mOS) in KPC mice compared to control groups (vehicles + isotype 12.5 days vs combo 41 days, p=0.0006). The 4-drug regimen is significantly better than single schedule arms (mOS: Adoi 19 days and AZD6738 + gemcitabine 13 days). A weekly ultrasound revealed a tendency for tumour stabilisation in mice treated with cytotoxic therapeutics plus adenosine inhibition, when compared to the control arm during the first 2 weeks of treatment (fig. 4K).

Combining adenosine blockade (Adoi) and immunotherapy (FCP) reduced significantly tumour growth of the IOT-resistant allograft model when compared to control treatment (65% tumour growth reduction; p=0.002) and FCP (35% tumour growth reduction; p=0.021) arms (suppl. fig. 6 C-D). Data from two separate experiments with the same controls showed that adding Adoi to FCP remains the best combination in controlling tumour growth (suppl. fig. 6E-G). These data suggest again that targeting the adenosine pathway in PDAC offers a new strategy to modulate the anti-tumour immune response.

### Targeting the adenosine pathway results in reprogramming of the TME in the IOT-resistant pancreatic cancer mouse model

To understand the role of adenosine modulation in reprogramming the TME, given the effect on tumour growth alone or in combination we analysed the changes in immune infiltration following anti-CD73 and AZD4635 treatment.

The TME of PDAC is an intricate structure that relies on the presence of multiple non-malignant cells. This TME is recapitulated in pre-clinical models of PDAC ^34^. In order to investigate the broader effect of Adoi in the 6419c5 PDAC model, given the effect we have seen on tumour growth when used alone or in combination, we performed a bulk RNAseq analysis of the 6419c5 model treated with Adoi or control (5 vs 6 mice, see methods). Following the 14-day treatment with AZD4635 and anti-CD73 the relative expression of 712 genes decreased by ≥50% (logfold <-0.58; fig. 5A). KEGG and GO Biological process pathway analysis revealed a dependency of the TME on the presence of extracellular adenosine (fig. 5B and suppl. fig. 7A). Genes associated with hypoxia response and vasculogenesis (e.g. *Hif1a*, *Slc2a1*, *Hilpda*, *Adm*, *Vegfa*, *Vegfd*), immunity and immune suppression (e.g. *Cd274*, *Cd209*, *Mrc1*, *Cd200*, *Il1a*, *Il6, Ptgs2*) and tumour stroma/ECM organisation (e.g. *Col5a3, Col6a3*, *Itga2*, *Mmp13, Mmp3*, *Mmp9*, *Ereg, Pthlh*) were significantly downregulated by the treatment (fig. 5B-C).

**Figure 5.**
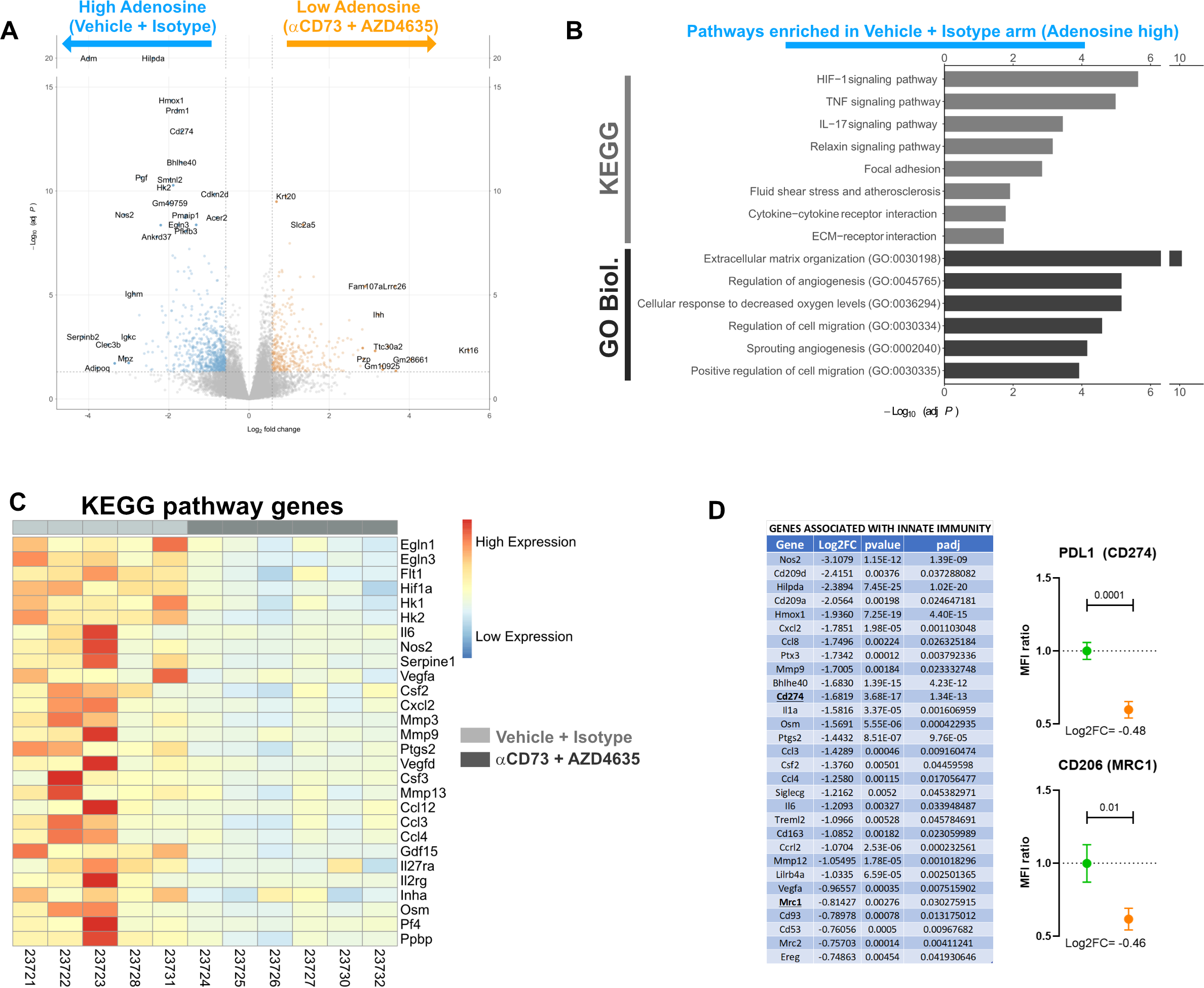
Adenosine inhibition remodels the PDAC tumour microenvironment. (A) Volcano plot related to 6419c5 allografts. Genes overexpressed in vehicle + isotype arm are on the left side (blue dots), and genes overexpressed in the anti-CD73 + AZD4635 arm are on the right side (Adoi, orange dots). (B) Enrichment bar plot of significantly overexpressed pathways in the vehicle + isotype group (adenosine high), according to KEGG and GO Biological processes. (C) Heatmap showing genes regulated during treatment (light grey for controls, dark grey for Adoi) which are part of the significantly different pathways according to KEGG. (D) Table (left) showing genes associated with innate immunity that are downregulated by Adoi treatment. Validation of RNAseq data (right graphs) through flow analysis of PD-L1 and CD206 expression (MFI ratio) on intra-tumoral live cells (8 mice in the control and 10 in the Adoi arms). The samples analysed for validation were from different mice from the ones used for RNAseq.

Of note, several functional and structural downregulated genes were associated with the innate immune system (fig. 5D left), particularly with M2 polarised macrophages (e.g. *Cd209*, *Mrc1*, *Mrc2*, *CD163*). RNAseq analysis also showed a significant decrease of the expression of *Cd274* (PD-L1) gene, suggesting a strong rationale for the use of adenosine inhibition as combination for immunotherapy studies involving immune checkpoint inhibitors. The downregulation of *Mrc1* (CD206) and *Cd274* (PD-L1) was validated analysing their protein expression on tumour-infiltrating live cells following Adoi treatment. As shown in fig 5D (right), the proteins of these genes were strongly downregulated in the adenosine inhibition arm (38% and 41% reduction of CD206 and PD-L1 on live cells respectively).

Furthermore, the adenosine signalling has previously been associated with the presence of hypoxia ^46, 47^; here we show for the first time that the presence of a functional extracellular adenosine pathway is responsible for the expression of several genes related to hypoxia (including *Hif1a*) suggesting the presence of a positive feedback loop.

Considering that the RNAseq analysis indicated that the innate immune system was affected by adenosine inhibition, we analysed the changes in immune infiltration following anti-CD73 and AZD4635 treatment. Flow cytometry analysis confirmed the findings highlighted by the RNAseq analysis. Following treatment, tumours were less likely to be infiltrated by M2 macrophages (median approximately 35000 vs 23000 cells/100mg; p=0.004; fig. 6 A-B), in particular PD-L1^+^ ones (median 79% vs 65%; p=0.004; fig. 6 B, right). IMC analysis revealed that the reduction of M2 macrophages either F4/80^+^ CD206^+^ (fig. 6C, E) or CD68^+^ CD206^+^ (fig. 6D, F) cells, was more prominent in viable tumour areas. Notably this reduced infiltration is present in areas other than those where adenosine is abundant, suggesting that adenosine is stimulating the production of factors affecting recruitment of macrophages in the viable tumour areas. Blocking the axis, also led to a reduced frequency of infiltrating Tregs (mean 42% vs 27%; p=0.03, suppl. fig. 7B-C) and a reduction of PD-L1 expression for all live cells within the tumour, more prominently CD45^+^ cells (p=0.008) as shown in suppl. fig. 7D. However, the expression of PD-L1 however declined in F4/80^+^ macrophages (p=0.02) but not in dendritic cells following treatment (suppl. fig. 7E). Of note, the combination of 2c5mIgG1 and AZD4635 is required to reduce the M2 macrophage infiltration (p=0.03, suppl. fig. 7F) and the ratio of M2/M1 macrophages (p=0.04; suppl. fig. 7G) in the tumour, while single agents fail to do so. Finally, IMC data also confirmed a reduction of total macrophages in the treated tumours as shown by flow, again more evident in the viable tumour regions (fig. 6C,D, F4/80 and CD68 panels and suppl. fig 7H). IMC also showed a global reduction of M1 macrophages (F4/80^+^ CD206^−^ MHCII^+^) (suppl. fig. 7 I-J) which was not apparent in the flow analysis. Further studies would further shed light on the relevance of these results, analysing the contribution of adenosine abundance and spatial distribution to modelling of tumour infiltration by distinct macrophage subpopulations beyond M1-M2 dichotomy.

**Figure 6.**
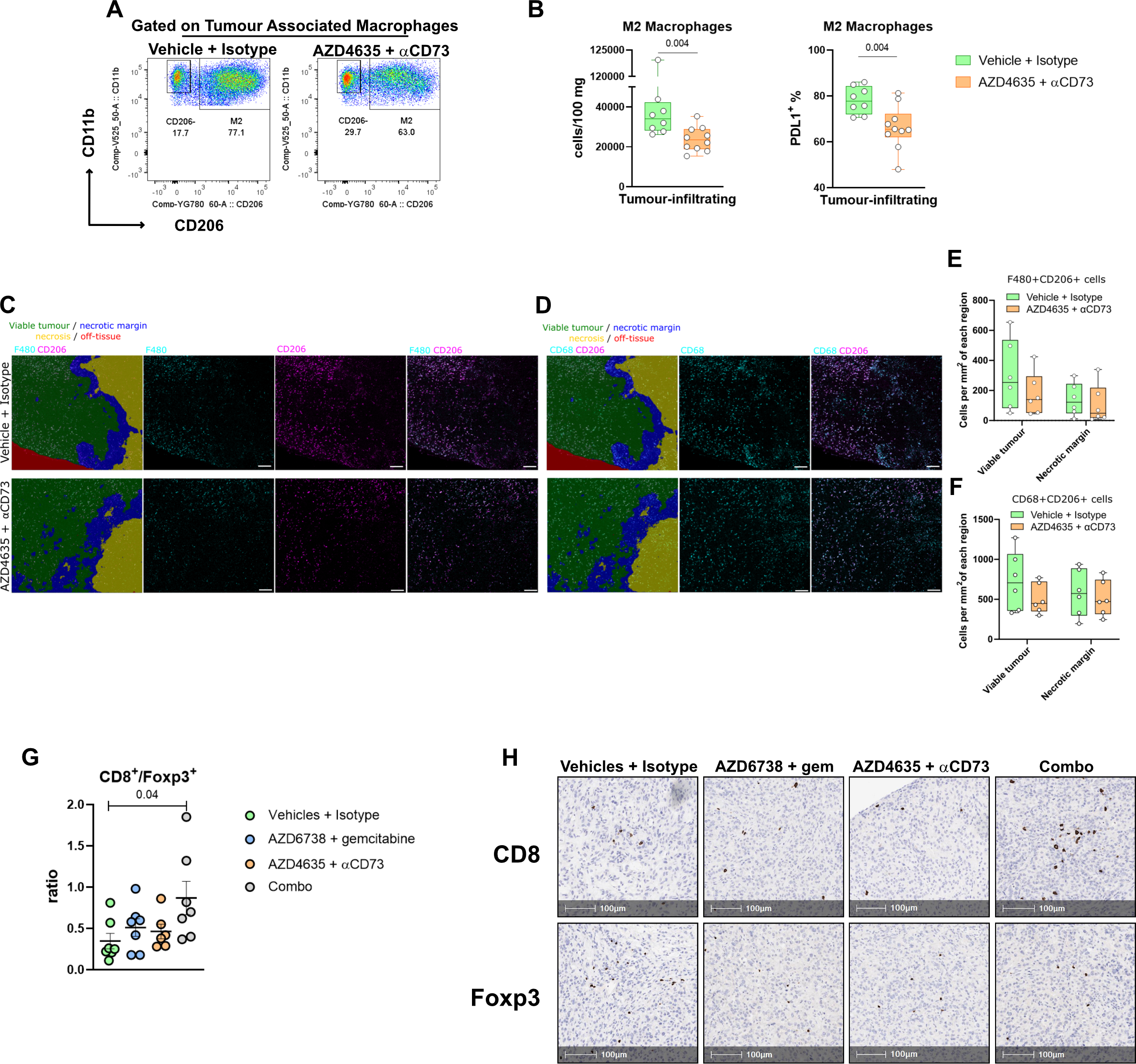
Immunosuppressive immune subpopulations are modulated following adenosine pathway blockade. (A) Representative flow cytometry plots showing tumour associated macrophages (TAM) infiltrating 6419c5 allografts following 14 day of treatment with vehicle + isotype (left panel) or anti-CD73 + AZD4635 (right panel). (B) M2 macrophage allograft infiltration (left, number of cells/100 mg of tumour) and (right) percentage of M2 macrophages positive for PD-L1 (8 and 10 mice per group analysed). (C-D) Representative IMC image of F4/80 and CD206 (C) and CD68 and CD206 (D) positive cells infiltrating a 6419c5 allograft merged (1^st^ panel of C and D) or not with segmentation following Adoi (lower panels) or control (upper). The tissue segmentation of the IMC image was performed by Random Forest Classification using all markers analysed. The scale bar on the IMC image is 200 μm. Segmentation shows viable tumour (green), necrosis (yellow), necrotic margin (blue) and off-tissue (red). (E-F) The bar plots show cell density (number of cells per mm2) of F480^+^CD206^+^ cells (E) and CD68^+^CD206^+^ (F) per segment area (6 mice per treatment group). (G-H) Immunohistochemistry analysis showing 6419c5 tumour-infiltrating CD8^+^/Foxp3^+^ ratio at day 14 of the following treatments: Vehicles + isotype (N=7), AZD6738 + gemcitabine (N=7), aCD73 + AZD4635 (N=6) and AZD6738 + gemcitabine + aCD73 + AZD4635 (N=7). (G) Representative IHC images of the latter. All data are represented as box and whisker plots. Statistical analysis was performed using Mann-Whitney test (B-H) and one-way ANOVA with post-hoc test analysis for multiple comparisons (E-H); p values are shown in the graphs when considered significant (p<0.05).

Considering the reduction of the immune suppression in the TME following Adoi, we explored changes associated with the adaptive immune system following combination of adenosine inhibition and cytotoxic or immunotherapy therapeutics. To assess whether the combination of Adoi plus ATRi/gemcitabine had an effect on the immune infiltration of the TME of IOT-resistant model (6419c5 allografts), we performed IHC staining for CD8 and Foxp3 and showed that the quadruple combination almost tripled the ratio CD8/Tregs (median 0.70 vs 0.25 of vehicles + isotype group; p=0.04; fig. 6G-H). The 4-drug combination produced an enhanced increase of the CD8^+^ cell infiltration (median 4.5 cells/mm^2^ vs 2.7 in the control group, 3.9 in the AZD6738/gem group and 2.8 in the AZD4635/αCD73 group) and reduction of Foxp3^+^ cell infiltration (median 6.5 cells/mm^2^ vs 8.1 in the control group, 9.3 in the AZD6738/gem group and 7.2 in the AZD4635/αCD73 group) (suppl. fig. 7K). Similarly, combination with IOT (Adoi + FCP) determined a further increase of the intra-tumoral CD8/Tregs ratio induced by FPC treatment (ratio means control 0.28 vs FCP/Adoi 6.10 p=0.008; suppl. fig. 7L) mostly due to the repression of the Treg recruitment induced by IOT alone.

### Adenosine-related gene expression is associated with phenotype and survival in human pancreatic cancer

To evaluate the importance of this pathway in the context of human PDAC and highlight the role in the formation of TME we scored our adenosine-related gene expression set against the PDAC subtypes published by Bailey ^39^. Of our 712 genes with a 50% decrease, 561 had a human ortholog and of these the z-scores of 517 genes were summed in each of the 97 patients with RNAseq data in Bailey et al. to obtain a score. We were able to show that the adenosine-related gene expression is mostly expressed in the aggressive squamous subtype (fig. 7A), that has been associated with a poorer prognosis. The score remained significantly higher even when comparing the squamous subtypes with the grouped non-squamous (fig. 7B).

**Figure 7.**
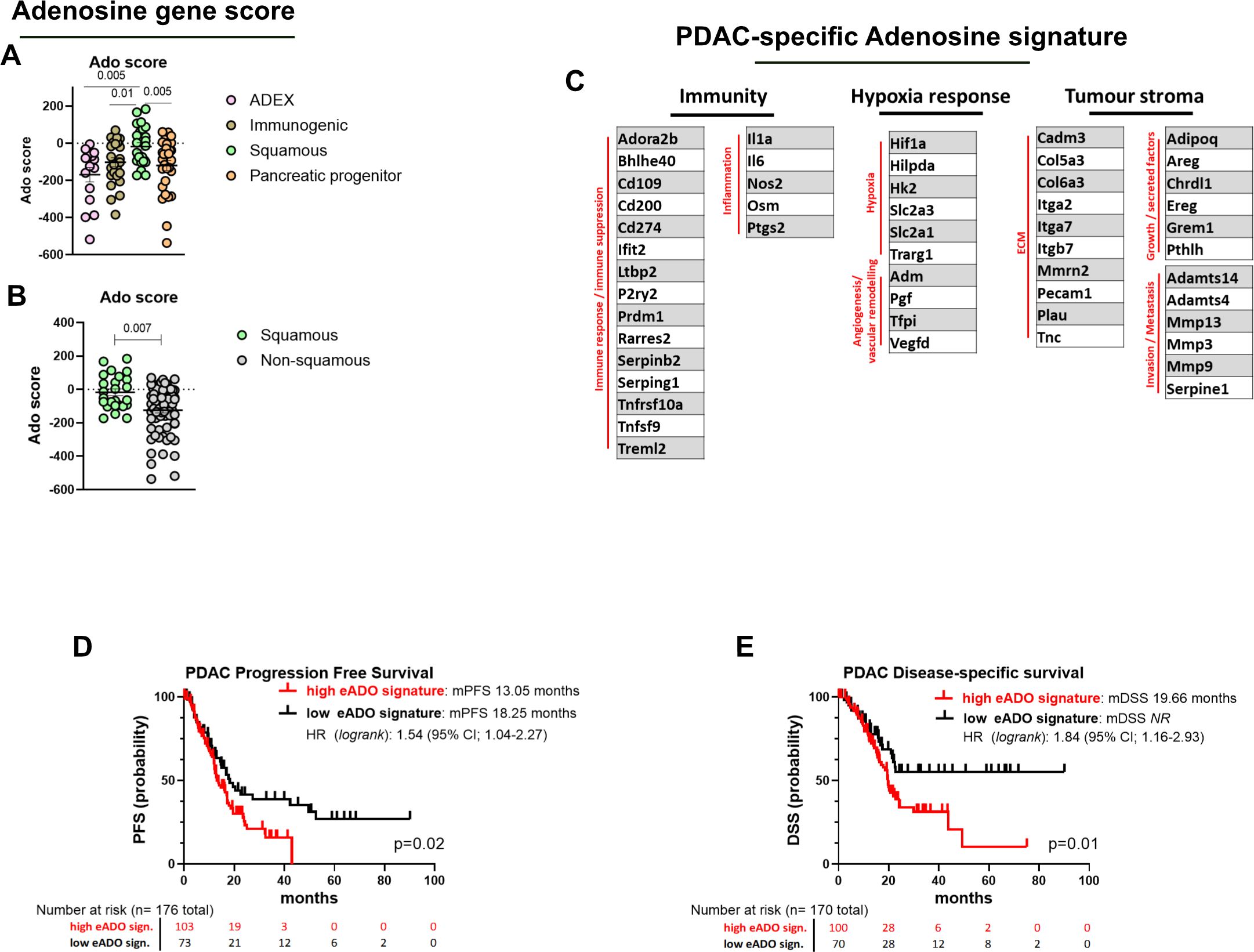
RNA gene expression demonstrates a crucial role for the adenosine pathway in correlation to human PDAC prognosis. (A-B) Human ortholog genes with a 50% downregulation following 14 days of Adoi, were scored using z-score derived from Bailey’s^39^ subtypes dataset. Genes scores comparing Adex, immunogenic, squamous and pancreatic progenitor (A) or squamous vs non-squamous (B) are shown. Each dot represents a single patient. (C) PDAC-specific gene signature of 52 genes used for analysis of the TCGA human PDAC dataset, related to the main pathways implied (immunity, hypoxia response and tumour stroma). (D-E) Gene signature applied to PDAC TCGA dataset (G) for progression free survival (PFS, 176 patients) and (H) disease specific survival (DSS, 170 patients) Kaplan-Meier. All data are presented as mean ± SEM when applicable. Statistical analysis was performed using one-way ANOVA with post-hoc test analysis for multiple comparisons (A), two-tailed unpaired Student’s t-test (B) and log-rank Mantel-Cox test to evaluate difference in survivals (D-E); p values are shown in the graphs when considered significant (p<0.05).

In order to create a PDAC-specific adenosine signature and to evaluate its performance in PDAC-specific outcomes, we created a signature using the 561 human orthologous genes with a ≥ 50% decrease in expression following Adoi treatment. Of these, 52 genes were selected for the signature (fig. 7C), that according to our RNAseq dataset were clearly associated with the pathway areas dependent on adenosine (hypoxia response, immunity, tumour stroma), were associated positively or negatively with PDAC prognosis, and were significantly co-expressed with CD73 and/or Adora2a in the PDAC genome datasets (TCGA or Bailey). Of the 176 and 170 patients with available progression free survival (PFS) and disease-specific survival (DSS) outcome respectively, we found that the presence of a high adenosine signature is associated with higher probability of PDAC progression (mPFS high Ado 13.05 vs low Ado 18.25, p=0.02; fig. 7D) and a poorer PDAC-specific survival (mDSS high Ado 19.66 vs low Ado *NR*, p=0.01; fig. 7E), suggesting again that the presence of a functional adenosine pathway has a detrimental role in human PDAC. These data should be considered in light of the fact that patients in the TCGA dataset are predominantly non-metastatic, and the presence of the adenosine signature seems to become relevant for PDAC associated death 20 months after diagnosis. The presence of a high adenosine signature appears to be associated with shorter disease-free-survival, (mDFS high Ado 23.5 months vs low Ado 49.7 months, p=0.12; suppl. fig. 7M).

## Discussion

Overall our data highlight that the generation of the extracellular adenosine is instrumental for the innate immune system in shaping a pro-tumorigenic, immune suppressive microenvironment in PDAC, in the context of a hypoxic milieu. We can speculate that this unfavourable environment may create the condition for a more aggressive PDAC phenotype which would then translate into the ability to escape the immune system, be resistant to cytotoxic treatment and metastasize readily.

Pancreatic ductal adenocarcinoma (PDAC) is projected to become the second highest cause of cancer-related death in the US within 10 years ^48^, and represents one of the major unmet needs of cancer treatment. Despite extensive efforts by laboratory and clinical scientists in the last 50 years, only 1% of patients diagnosed with PDAC today will survive for 10 years (https://www.cancerresearchuk.org/health-professional/cancer-statistics-for-the-uk). The response rate following standard treatments is poor, usually short-lasting, and associated with significant treatment related toxicity ^1^. In the past few years, immunotherapy has provided new hope in the treatment of several types of cancer, and has dramatically changed the life expectancy of many patients with metastatic disease ^49^. However, this has not been true for patients with PDAC which is associated with a very low response rate to immunotherapy, usually confined to MSI-H/dMMR tumours, found only rarely in this disease ^2^.

The role of the innate immune system in the generation of an immune suppressive/pro-tumorigenic microenvironment in PDAC is well known. The presence of marked infiltration of macrophages has been identified as an independent predictive factor of the aggressiveness and prognosis of PDAC, in patients ^23, 24^. Only recently, three phase I clinical trials in patients with PDAC have shown that targeting the innate immune system can have impact in patients with PDAC. A phase I trial published on Lancet Oncology, showed that a combination of anti PD-1 and CD40 agonist, added to gemcitabine and nab-paclitaxel, led to 60% of response rate, with some durable responses ^50^. In addition, the inhibition of the CXCL12/CXCR4 axis has been demonstrated to modify the immunosuppressive TME of PDAC and CRC patients ^51, 52^.

The extracellular adenosine pathway has also been shown to influence the TME fostering the immune suppression provided by some innate immune subpopulations (as myeloid and NKs) and inhibiting the function of the adaptive immune system, in particular, T-cells ^6, 33, 44^. By activating its receptors, eAdo is able to increase the intracellular concentration of cAMP which leads to the induction of a M2 phenotype of macrophages and block the secretion of IL1β increasing the release of CXCL1, IL-6, IL-10 and IL-8 among others from myeloid population which are known to orchestrate immune exclusion ^8, 53^. eAdo also favours the formation and maintenance of Treg cells ^8^, which are known to favour cancer progression and IOT resistance.

A recent publication, shows that genomic targeting on mouse PDAC cells of CD73 leads to a reduced in vivo tumour formation and change in the circulating and infiltrating immune system^54^.

However, to date, little was known about the expression of the adenosine pathway in the context of the innate and adaptive immune system in PDAC, how the extracellular adenosine is generated and what are the targets of adenosine also in regard to its spatial distribution and formation of adenosine.

Our results show for the first time, that the mechanism of generation of extracellular adenosine in pancreatic cancer TME is finely orchestrated by tumour infiltrating myeloid cells and tumour cells, due to the expression of high level of CD39 in infiltrating myeloid cells and CD73 on both cell types. We have also demonstrated that the pathway can be overexpressed in T-cells infiltrating the tumours, regardless of their activation status (fig. 1,2 and suppl. fig. 2,4). The distribution of extracellular adenosine is spatially heterogeneous and a high level of extracellular adenosine correlates with the presence of a hypoxic environment and is favoured by the presence of necrosis, where the myeloid population is enriched (fig.3). Necrosis is common in human PDAC and related to poor prognosis for all stages ^55^. The enrichment of a CD39^+^ CD73^+^ double population, potentially able to independently produce adenosine, does not seem to correlate with IOT-resistant or responsive tumour models, but there is a difference when the target of adenosine (Adora2a receptor), is considered. Adora2a on myeloid populations, in particular in pro-tumorigenic M2 macrophages but also in antigen presenting cells, is differentially expressed in regard of IOT response phenotype, with the resistant tumours abundantly overexpressing the receptor in these populations (fig.3 and suppl. fig. 4). The bulk RNAseq analysis of tumour treated with adenosine inhibition revealed indeed a broader role for adenosine in PDAC TME (fig. 5). Genes related to immunosuppression and innate immunity recruitment (*Cd274*, *Csf2*, *Cxcl2*, *Ccl3*, *Ccl4*, *Ccl12*, *Il1a*, *Osm*, *Il6*), angiogenesis (*Vegfa*, *Vegfd*, *Adm2*, *Flt1*, *Pgf*, *Egln1*) and cell-ECM interaction (*Adam19*, *Adamts14*, *Adamts4*, *Adamts5*, *Col5a3*, *Col6a*, *Itga2*, *Itga7*, *Mmp3*, *Mmp9*, *Mmp12*) indicated the targeting of population of cells responsible for the acquisition of a pro-tumorigenic, pro-metastatic, pro-fibrotic and immune resistant phenotype. Further, the downregulation of genes associated with hypoxia (*Hif1a*, *Hilpda*, *Nos2*, *Hk1*, *Hk2*, *Egln3*) following treatment shows not only that the adenosine pathway is induced during hypoxia (e.g. CD73 and Adora2b), but that the hypoxia response is also dependent on the presence of the adenosine pathway in what we can speculate is a positive feedback loop. The inhibition of adenosine led to a reduced infiltration of M2 macrophages further from the hypoxic regions where adenosine is most abundant, suggesting that the effect of adenosine is stimulating the secretion of factors that are recruiting monocytes into the tumour microenvironment, replenishing macrophage infiltration (fig. 6 and suppl. fig. 7). Notably, targeting the pathway can reduce tumour growth in an IOT-resistant model, improving the response to cytotoxic and immunotherapy combinations (fig. 4 and suppl. fig. 6) even in historically therapy-resistant models as the KPC mice.

Targeting adenosine would represent an alternative strategy to reduce the infiltration of pro-tumorigenic macrophages in PDAC lesions. One approach has been the administration of a CSF1R inhibitor ^32^, which has shown promising pre-clinical data, that have not been translated in human so far. A recent publication shows that in CRC mouse models, the use of anti CSF1R treatment spares a subpopulation of macrophages characterised by the expression of *Cd274* (PD-L1), *Vegfa*, *Hilpda*, *Bhlhe40*, *Mmp12*, *Cebpb*, *Hmox1* among others ^28^. Given that all of these genes are among the immune suppressive and vasculogenic molecules that seem to be strongly downregulated by adenosine inhibition and our data show a reduction of some subpopulation of PD-L1^+^ macrophages, we can speculate that adenosine inhibition could potentially target these populations of pro-angiogenic, immune suppressive macrophages.

Information provided by the PDAC specific adenosine signature, indicate that the adenosine pathway appears to play a role in progression and survival of human PDAC, due to the ability of the adenosine inhibition to profoundly reprogram the tumour microenvironment in PDAC models (fig. 5–6 and suppl. fig. 7). Gathering more information on the role of this pathway in human cancers should be a priority, retrospectively evaluating and then prospectively stratifying the patients on the basis of histopathological/radiological features and the spatial distribution of adenosine.

In summary, we have shown for the first time that tumour-infiltrating myeloid immune cells contribute to the generation of extracellular adenosine in the context of PDAC, in correlation with the presence of hypoxia. Macrophages in particular, express high levels of Adora2a receptor in PDAC models and targeting the adenosine/myeloid axis remodels the TME. Data from IMC, flow cytometry and RNAseq suggest that the adenosine pathway is fundamental to the formation of a pro-tumorigenic, immunosuppressive TME, and its expression is associated with an aggressive phenotype and poor survival in human PDAC. The re-modelling of the TME caused by the inhibition of the adenosine pathway, translate into a delay of PDAC tumour growth. Finally, targeting extracellular adenosine should be considered as an alternative to improve the efficacy of cytotoxic and IOT in future PDAC-specific clinical trials.

## Author contributions

V.G. and D.I.J conceptualised the project. V.G., A.D., H.H., K.W., J.P.M., J.E. and DIJ designed the experiments. V.G., A.D., H.H., K.W., H.B., S.A.K., S.Y.L., B.P.K. conducted the experiments. V.G., A.D., H.H., K.W., S.I., F.M.R., R.B., J.P.M, S.J.D. J.E. and D.I.J analysed the data. J.E.D.T. R.G., J.P.M., S.J.D., A.G.S, J.E., D.I.J. provided supervision and V.G., A.D., H.H., K.W., S.I., J.P.M., S.J.D., A.G.S, J.E., D.I.J. discussed the results. V.G. and D.I.J. wrote the manuscript. All authors read, review, edited and approved the final manuscript.

## Acknowledgments

All CRUK CI authors received research funding from Cancer Research UK (Nos. C14303/A17197 and C9545/A29580). The Li Ka Shing Centre where this work was performed was generously funded by CK Hutchison Holdings Limited, the University of Cambridge, CRUK, The Atlantic Philanthropies and others. This work was supported by Cancer Research UK (C9685/A27444) funding to VG. The authors wish to thank all the CRUK Cambridge Institute core facilities, in particular Research Instrumentation & Cell Services (RICS), Flow Cytometry, Histopathology, Biological Resource Unit (BRU), Genomics and Genome editing cores. VG would like to thank Maike de la Roche (CRUK Cambridge Institute - University of Cambridge) and Klaus Okkenhaug (Department of Pathology - University of Cambridge) for helpful suggestions and advices. This study was also supported by Cancer Research UK Precision Panc grant C96/A25238. HB holds a PhD studentship at the University of Cambridge which is supported jointly by the University of Cambridge Experimental Medicine Training Initiative (EMI) programme in partnership with AstraZeneca (EMI-AZ) and the NIHR Biomedical Research Centre and SYL is founded by the Cambridge Trust (Cambridge International Scholarship). Work by JT / KW is funded by a core grant award from the MRC (MC_UU_00025/12). S.A.K. and. J.P.M. were supported by Cancer Research UK core funding to the Beatson Institute (A17196 and A31287) and to J.P.M. Lab (A29996). The KPCY-derived cell lines were a kind gift of Ben Stanger (University of Pennsylvania). AZD4635, AZD6738 and the antibodies anti-CD73 (2C5, murine IgG1-Fc), anti-PD-L1 (AB740080 D265A), NIP228 muIgG1 isotype, NIP228 muIgG1 D265A isotype, were kindly provided by AstraZeneca. The results shown here are in part based upon data generated by the TCGA Research Network: https://www.cancer.gov/tcga.

## Notes

Financial support All CRUK CI authors received research funding from Cancer Research UK (Nos. C14303/A17197 and C9545/A29580). The Li Ka Shing Centre where this work was performed was generously funded by CK Hutchison Holdings Limited, the University of Cambridge, CRUK, The Atlantic Philanthropies and others. This work was supported by Cancer Research UK (C9685/A27444) funding for VG. This study was also funded by Cancer Research UK Precision Panc grant C96/A25238. HB is funded by the University of Cambridge Experimental Medicine Training Initiative programme in partnership with AstraZeneca (EMI-AZ) and SYL is funded by the Cambridge Trust (Cambridge International Scholarship). Work by JT and KW is funded by a core grant award from the MRC (MC_UU_00025/12). S.A.K. and. J.P.M. were supported by Cancer Research UK core funding to the Beatson Institute (A17196 and A31287) and to J.P.M. Lab (A29996).

### Competing Interest Statement

AD, HH, BPK, FMR, RG, SJD, AGS and JE are Astrazeneca employees and own company stocks and shares. HB holds a studentship partially funded by AstraZeneca.

### Summary of Updates

- A new experiment in KPC mouse model of pancreatic cancer has been added. This further supports the conclusion of the manuscript. - RNAseq validation

