## Supplementary Figure 1 to 7 for "Defining the spatial distribution of extracellular adenosine revealed a myeloid-dependent immunosuppressive microenvironment in pancreatic ductal adenocarcinoma"

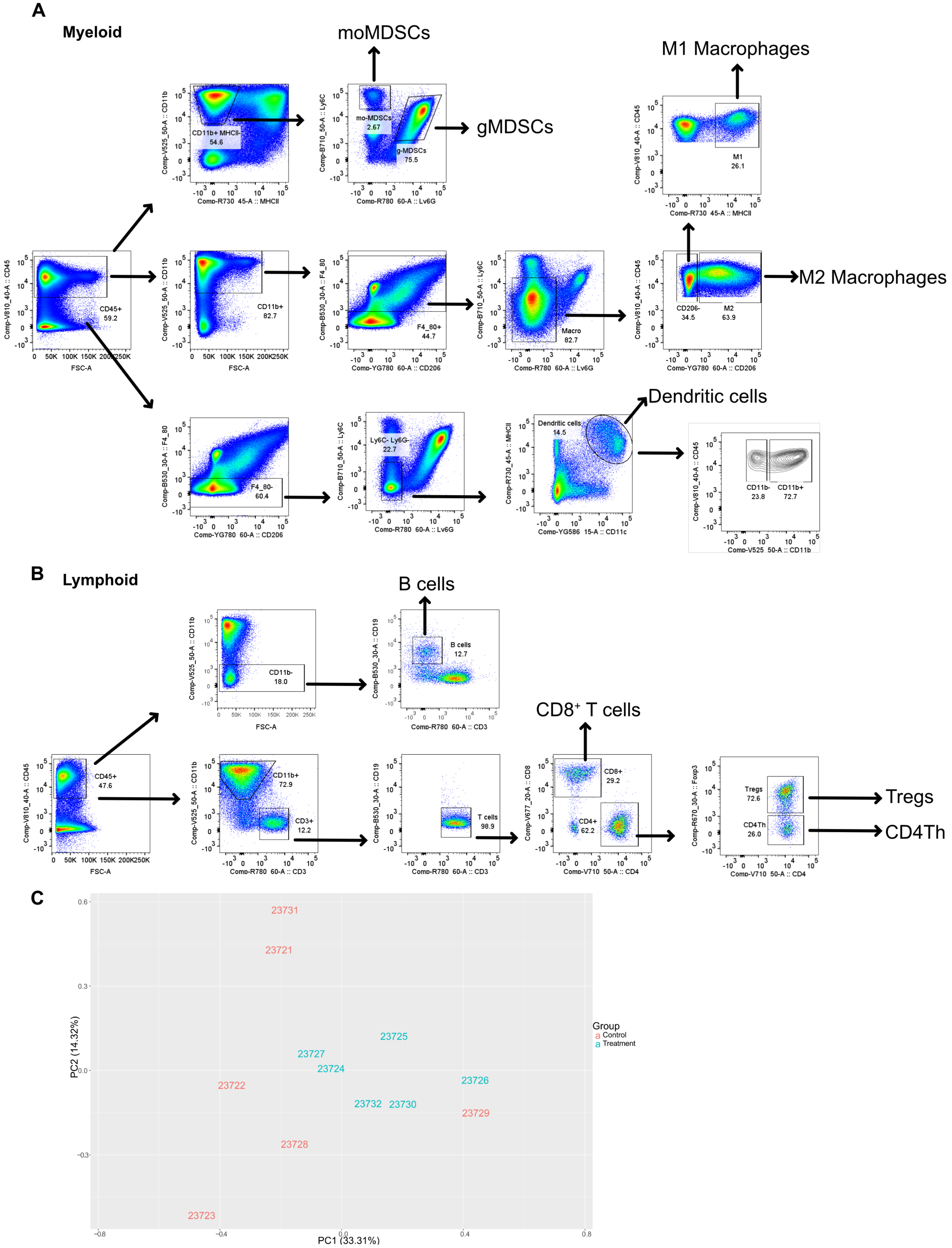


**Supplementary figure 1. Gating strategy and RNAseq PCA.** Flow cytometry dot plots showing gating strategy for myeloid (A) and lymphoid (B) immune populations. (C) PCA analysis of RNAseq related to mice with 6419c5 allografts treated with vehicle+isotype (n=6) or αCD73+AZD4635 (n=6) for 14 days.

### Supplementary figure 2

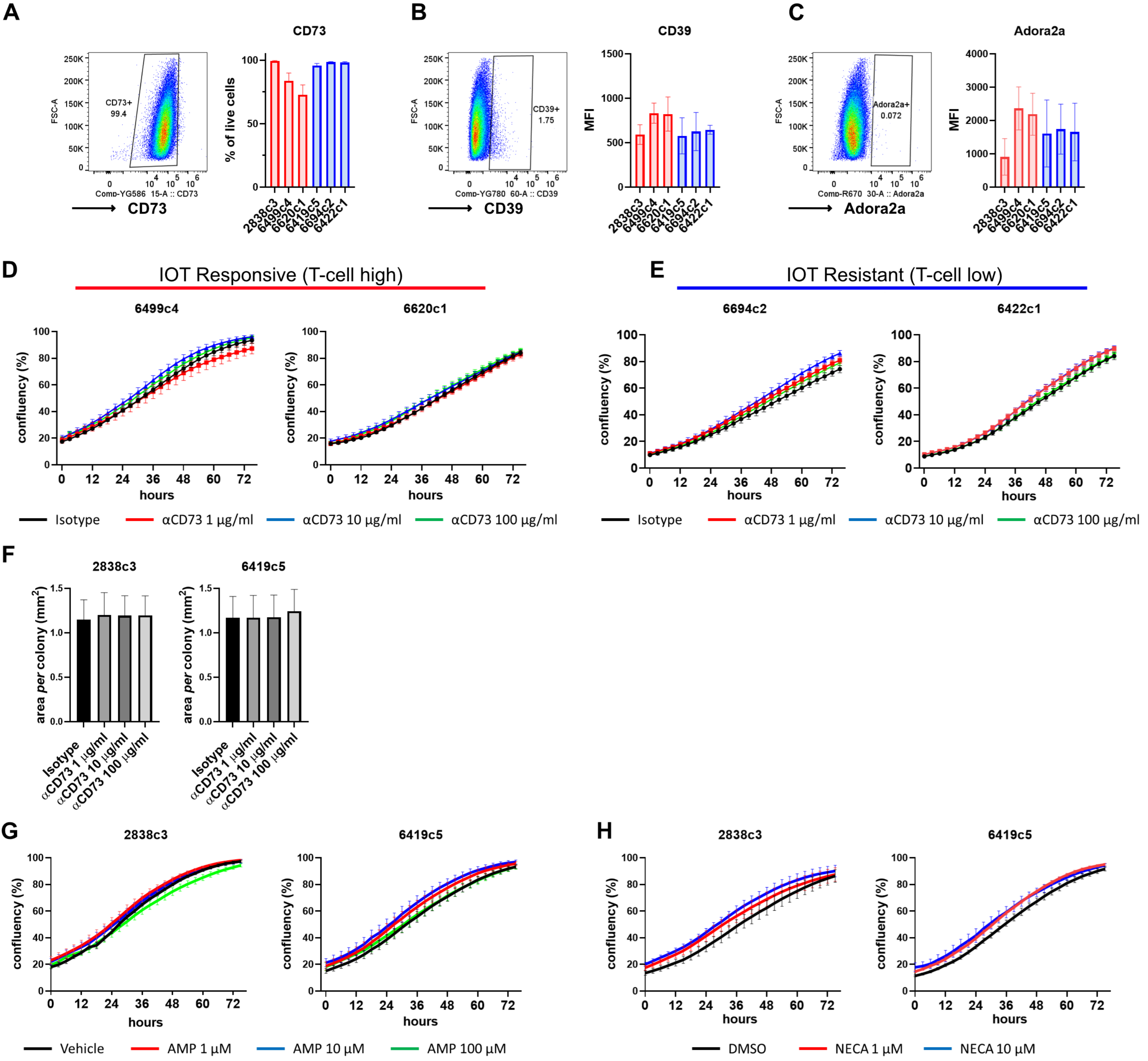


**Supplementary figure 2. Expression of the adenosine pathway on KPC-derived cell lines and response to anti-CD73 antibody.** (A-C) Representative dot plots for CD73 (A), CD39 (B) and Adora2a (C) expression on KPC-derived cell lines and associated bar graphs with % of positive cells per cell line. (D-E) Confluency (%) of KPC-derived cell lines upon anti-CD73 in vitro treatment up to 72 hours. Concentrations are shown below graphs. (F) Cell colony area in mm² following 8-day treatment with increasing concentrations of anti-CD73 and isotype (100 µg/ml) for 2838c3 (left) and 6419c5 (right). (G-H) Confluency (%) of 2838c3 (left) and 6419c5 (right) upon 72-hour in vitro treatment with increasing concentrations of AMP (G) and NECA (H). Concentrations are shown below the graphs. All experiments have been repeated at least 3 times, with at least 3 technical replicates each time. All data are presented as mean +/- SEM. Statistical analysis was performed using one-way ANOVA with post-hoc test analysis for multiple comparisons (A-C, F) and mixed-effect models (D-E, G-H); p values are shown in the graphs when significant ($p<0.05$).

**Supplementary figure 3**

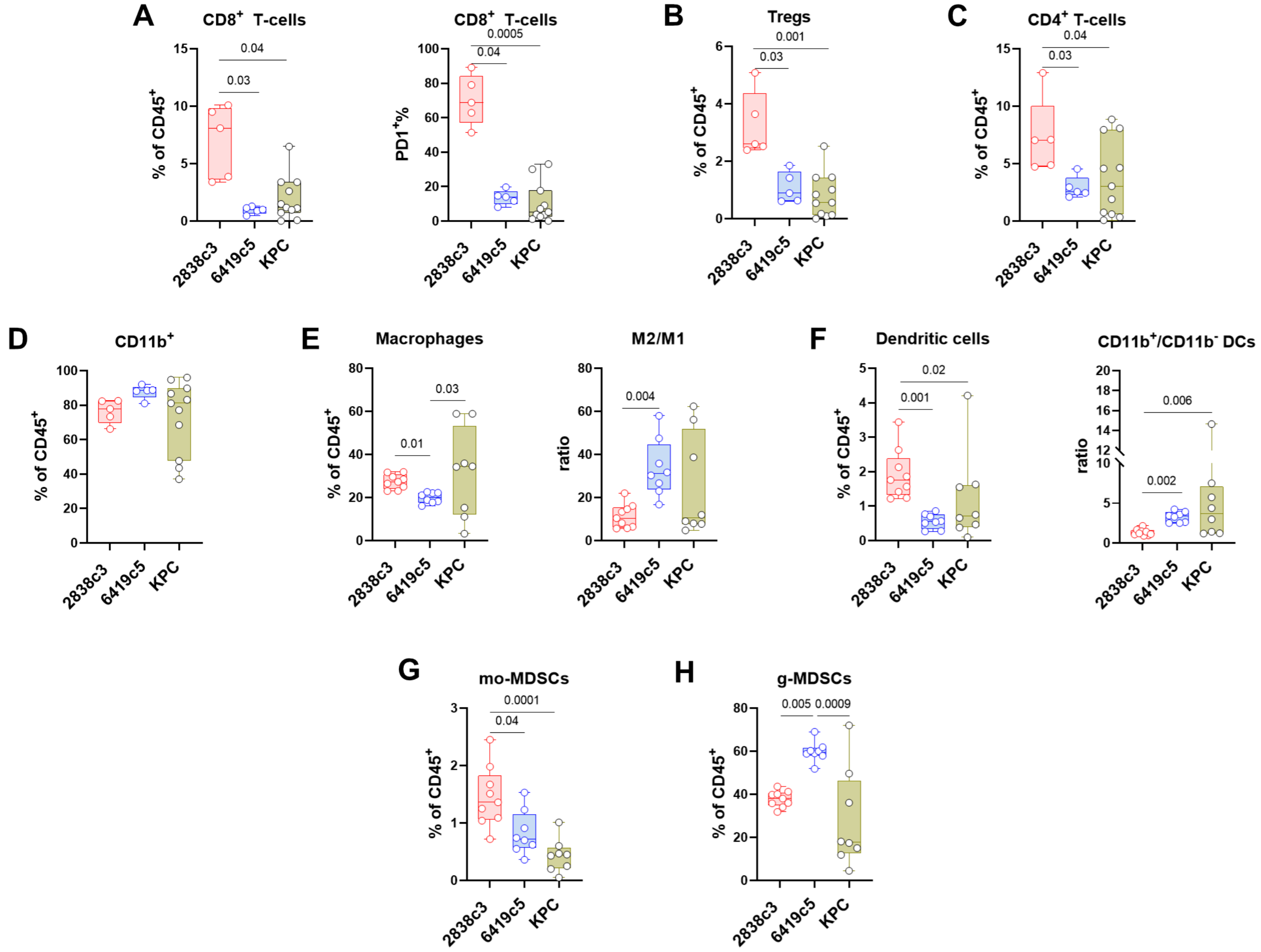


**Supplementary figure 3. KPC tumour immune infiltration has similarity to the IO resistant derived subcutaneous allografts.** (A-H) Flow analysis of tumour-infiltrating immune subpopulations for 2838c3 and 6419c5 s.c. allografts and KPC autochthonous tumours. The following are shown: (A) % of CD8 T-cells out of total CD45+ cells (left) and % of PD1+ cells out of total CD8 T-cells (right). Percentage out of total CD45+ tumour-infiltrating cells for Tregs (B), CD4 T-cells (non Tregs, C), CD11b+ cells (D), total macrophages (E, and M2/M1 macrophages ratio, right), dendritic cells (F, and CD11b+/CD11b- dendritic cells ratio, right), mo-MDSCs (G) and g-MDSCs (H). Box and whisker plots are presented. Statistical analysis was performed using one-way Anova with post-hoc test analysis for multiple comparisons; p values are shown in the graphs when significant ($p<0.05$).

### Supplementary figure 4

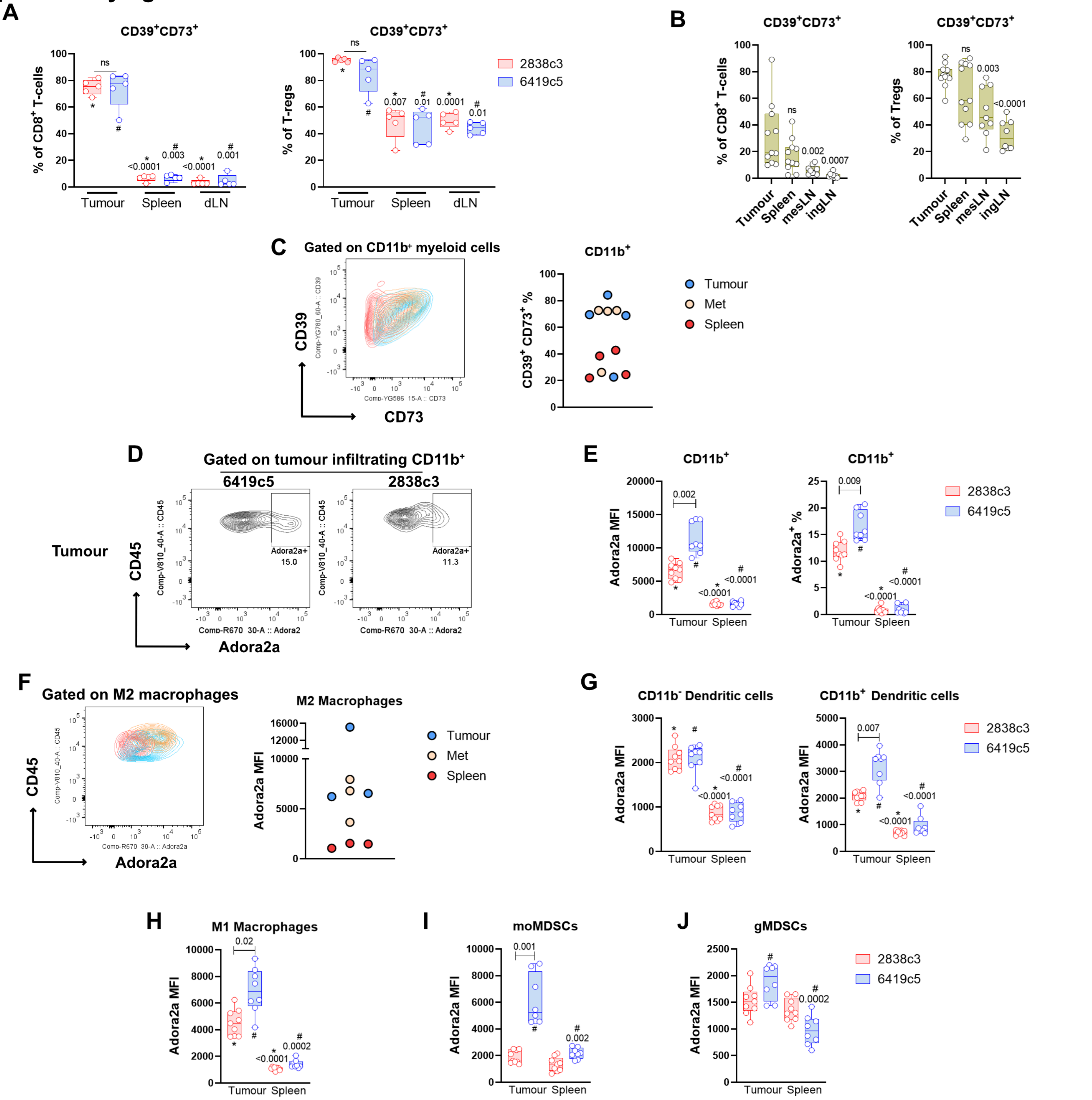


**Supplementary figure 4. Immune cells infiltrating PDAC models double express CD39 and CD73 and Adora2a receptor expression in increased in myeloid cells, in particular M2 macrophages.** (A) CD39 and CD73 expression on CD8 T-cells (left) and Tregs (right) for 2838c3 and 6419c5 allografts, compared to matched spleens and draining lymph nodes. (B) CD39 and CD73 expression on CD8 T-cells and Tregs in KPC mice bearing pancreatic tumours. Comparison matched with spleens, mesenteric lymph nodes and inguinal lymph nodes. (C) Contour plot (left) overlapping CD39 and CD73 expression on CD11b$^+$ cells from tumour (blue), spleen (red) and metastatic liver lesion (orange) of a single KPC mouse. Percentage of CD11b$^+$ cells double expressing CD39 and CD73 in tumours (blue), spleens (red) and metastases (orange) of KPC mice. (D) Plot of Adora2a expression on CD11b$^+$ cells infiltrating 6419c5 (left) and 2838c3 (right) allografts. (E) Adora2a expression as MFI in CD11b$^+$ cells (left) or percentage out of total Cd11b$^+$ infiltrating PDAC allografts (right). (F) Contour plot (left) overlapping Adora2a expression on M2 macrophages from tumour (blue), spleen (red) and metastatic liver lesion (orange) of a single KPC mouse. Adora2a expression (MFI) of M2 macrophages in tumours (blue), spleens (red) and metastases (orange) of KPC mice. (G-J) Adora2a expression (MFI) in (G) CD11b$^-$ (left) and CD11b$^+$ (right) dendritic cells, M1 macrophages (H), mo-MDSCs (I) and gMDSCs (J). Box and whiskers are presented for all the graphs. Statistical analysis was performed using one-way Anova with post-hoc test analysis for multiple comparisons; p values are shown in the graphs when significant ($p<0.05$). Compared groups are connected by lines or have the same symbol # or *. For (B) p values shown are representative of comparison to tumour group.

### Supplementary figure 5

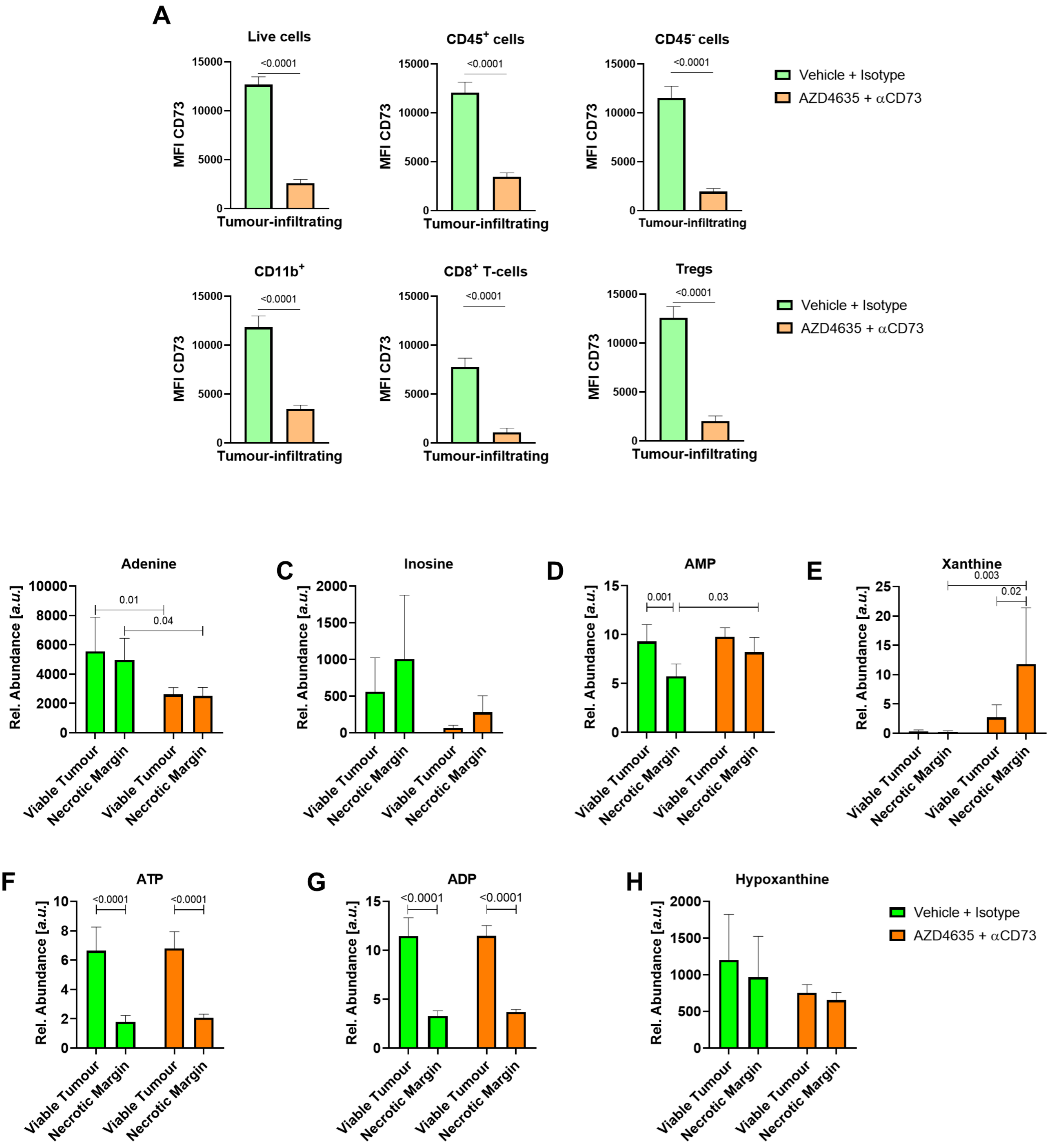


**Supplementary figure 5. Effect of adenosine inhibition on CD73 expression and tumour metabolites.** (A) Flow analysis of CD73 expression (MFI) following 14-day treatment with αCD73 + AZD4635 (N=10) or vehicle + isotype (N=8) in total live cells (upper left), total CD45+ cells (upper middle), total CD45- (upper right), CD11b+ cells (lower left), $CD8^+$ T-cells (lower middle) and Tregs (lower right). (B-H) Relative abundance of Adenine (B), Inosine (C), AMP (D), Xanthine (E), ATP (F) , ADP (G) and Hypoxanthine (H) in 6419c5 allografts treated with vehicle + isotype or αCD73 + AZD4635 analysed through MSI. All data show mean +/- SEM. Statistical analysis was performed using two-tailed unpaired Student's t-test (A) and one-way ANOVA with post-hoc test analysis for multiple comparisons (B); p values are shown in the graphs when significant ($p<0.05$).

### Supplementary figure 6

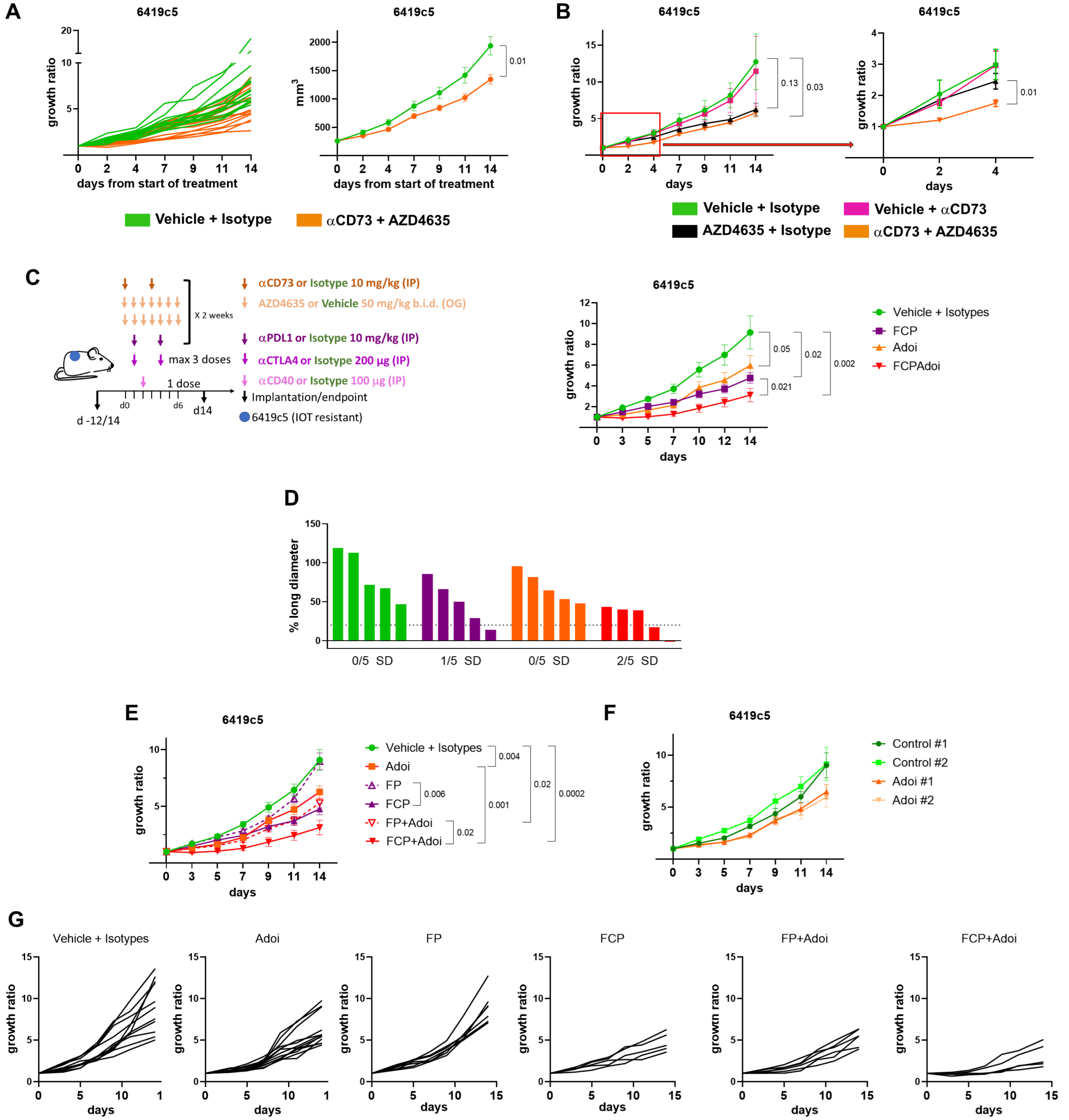


**Supplementary figure 6. Inhibiting the adenosine pathway reduces tumour growth in vivo, and improves tumour control when combined with cytotoxic or immunotherapy.** (A) Tumour growth ratio per single mouse (left) and tumour volume (in mm³) for mice treated with vehicle + isotype (N=15) or αCD73 + AZD4635 (N=16). (B) Tumour growth ratio of 6419c5 allografts treated with AZD4635 (N=5) and αCD73 (N=5) or combination (N=6) of the two (and compared to vehicle + isotype, N=5), only the first 4 days of treatment (right graph). (C) Schedule (left) and tumour growth ratio (right) of 6419c5 tumour allografts treated as following (N=5 mice per group): vehicle + isotypes, FCP (anti-CD40 agonist, anti-CTLA4, anti PD-L1), anti-CD73 + AZD4635 (Adoi), FPCAdoi. (D) Percentage change in the long diameter length following 14 days of treatment per group. Number of mice with stable disease (SD, <20% increase and <30% decrease) are shown at the bottom. (E) Tumour growth ratio of two merged experiments having similar control and Adoi arms growth in F. The following arms were analysed: Vehicle + Isotypes (N=11, control #1 N=6, control #2 N=5), αCD73 + AZD4635 (Adoi N=12, Adoi#1 N=7, Adoi#2 N=5), αCD40 agonist + αPDL1 (FP, N=7), αCD40 agonist + αPDL1 + αCTLA4 (FCP, N=5), FP + Adoi (N=6), FCP + Adoi (N=5). (G) Individual mouse tumour growth ratio for each treatment arm shown in D. Data show mean +/- SEM. Statistical analysis was performed using mixed-effects model (A-C, E); p values are shown in the graphs when significant ($p<0.05$).

### Supplementary figure 7

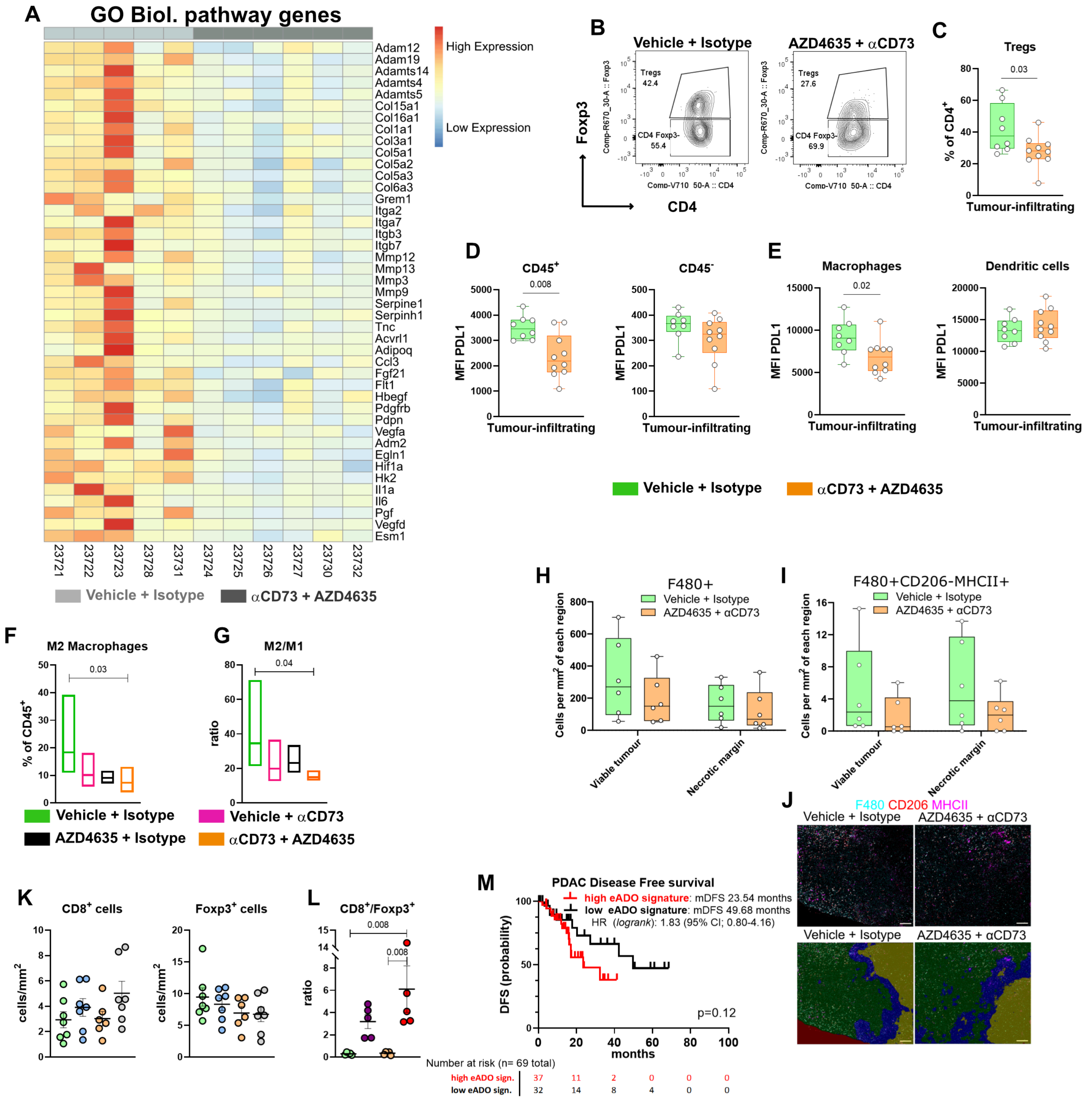


**Supplementary figure 7. TME modulation following adenosine inhibition and adenosine signature in human PDAC.** (A) Heatmap showing genes regulated during treatment (light grey for controls, dark grey for Adoi) which are part of the significantly different pathways according to GO Biological process. Some of the genes are also part of the Heatmap in fig. 5A. (B) Representative flow cytometry plot showing regulatory T-cells (Tregs) infiltrating 6419c5 allografts following 14 days of treatment with vehicle + isotype (left panel) or anti-CD73 + AZD4635 (right panel). (C) Infiltration of allografts by Tregs (number of cells/100mg of tumour; 8 and 10 mice per group analysed). (D-E) Flow analysis of PDL1 expression in $CD45^+$ cells (D, left), $CD45^-$ cells (right) and macrophages (E, left), dendritic cells (right) following 14-day treatment of vehicle + isotype (Control) or anti-CD73 +AZD4635 (Adoi). (F-G) Flow analysis on the effect on the TME of the treatments shown in suppl. fig. 6B, representing M2 macrophages tumour infiltration (F) and M2/M1 ratio (G). (H-J) IMC analysis for the spatial distribution (viable tumour and necrotic margin) of 6419c5 allografts in C57Bl/6 mice treated or not with Control or Adoi for 14 days. $F4/80^+$ cell (H) and M1 macrophages ($F4/80^+$ $CD206^-$ $MHCII^+$, I-J) are shown. (K) 6419c5 tumour-infiltrating $CD8^+$ (left) and $Foxp3^+$ (right) following ATRi/gem and Adoi experiment in fig.6G (L) Immunohistochemistry analysis following IOT and Adoi experiment in suppl. fig. 6C showing the ratio $CD8^+/Foxp3^+$ cells. (M) Human PDAC disease free survival based on 52-gene adenosine signature used for analysis of TCGA dataset (N=69 patients). Statistical analysis was performed using Mann-Whitney test (C-E), one-way Anova (F-G) with post-hoc test analysis for multiple comparisons (H-I,K-L), and log-rank Mantel-Cox test (L); p values are shown in the graphs when significant ($p<0.05$).
