## Supplementary Methods and Materials for "Defining the spatial distribution of extracellular adenosine revealed a myeloid-dependent immunosuppressive microenvironment in pancreatic ductal adenocarcinoma"

### Supplementary Materials and Methods

#### Immunohistochemistry (IHC)

The specific antibody details and conditions are listed in the table below.

| Target | Catalogue No. | Dilution/Conc. | Retrieval | Modifications |
|---|---|---|---|---|
| CD8 | Cell Signaling Technology, 98941 | 1:200 | Tris EDTA, 20’ | Protein Block |
| FOXP3 | Affymetrix, 14-5773 | 5 ug/ml | Tris EDTA, 20’ | Protein Block & Anti-rat secondary |
| p53 | Novocastra, NCL-L-p53-CM5p | 1:500 | Tris EDTA, 20’ | Protein Block |

#### Bioinformatics analysis: References

edgeR: (1)

DESEq2: (2)

Enhanced Volcano: Blighe K, Rana S, Lewis M (2020). EnhancedVolcano: Publication-ready volcano plots with enhanced colouring and labeling. R package version 1.8.0, https://github.com/kevinblighe/EnhancedVolcano.

Enrichr: (3)

Pheatmap: Raivo Kolde (2019). pheatmap: Pretty Heatmaps. R package version 1.0.12. https://CRAN.R-project.org/package=pheatmap

Scater/scuttle: (4)

Hisat: (5)

Samtools: H. Li, B. Handsaker, A. Wysoker, T. Fennell, J. Ruan, N. Homer, G. Marth, G. Abecasis, R. Durbin, 1000 Genome Project Data Processing Subgroup, The Sequence alignment/map (SAM) format and SAMtools, Bioinformatics, 25 (2009), pp. 2078-2079, 10.1093/bioinformatics/btp352

**Analysis of human PDAC available datasets and generation of PDAC-specific adenosine signature: References**

Bailey's subgroup analysis adenosine scoring and adenosine signature**:** (6)

**Mass spectrometry Imaging (MSI)**

Data segmentation pipeline applied to the MSI data: a) Co-registration of MSI data and the post-analysis H&E stained tissue sections. b) Annotation of the MSI data based on the underlying histology to generate a PLS-DA based machine learning classifier. c) Evaluation of the accuracy of the raised classifier based on 10-fold cross validation within the training and validation datasets. A minimum accuracy threshold of 95% in the training data and 90 % for the validation data was applied for all classification of MSI data throughout this work. d) Classification map as derived from the pixel-wise classification based on an accepted classifier overlayed with the H&E stained tissue section. e) Data reduction via principal component analysis (PCA) to visually identify components identifying variance in a spatial segment (in this case the necrosis adjacent tissue compartment). f) The top loadings of the components highlighting the area of interest were used to generate a feature list to perform unsupervised segmentation using bisecting k-means clustering on the pixel belonging to the tumour tissue. The yellow cluster corresponds to the necrotic margin, the blue cluster to viable tumour and red cluster was excluded as it contains pixel in the transition between viable tumour and necrotic margin and other pixel with an intrinsic high variance. g) Scoring plots of the PCA for the components displayed in e) and coloured corresponding to the cluster they were assigned to during the unsupervised segmentation in f). We ran the output by a trained pathologist for confirmation of the overall accuracy of the approach.

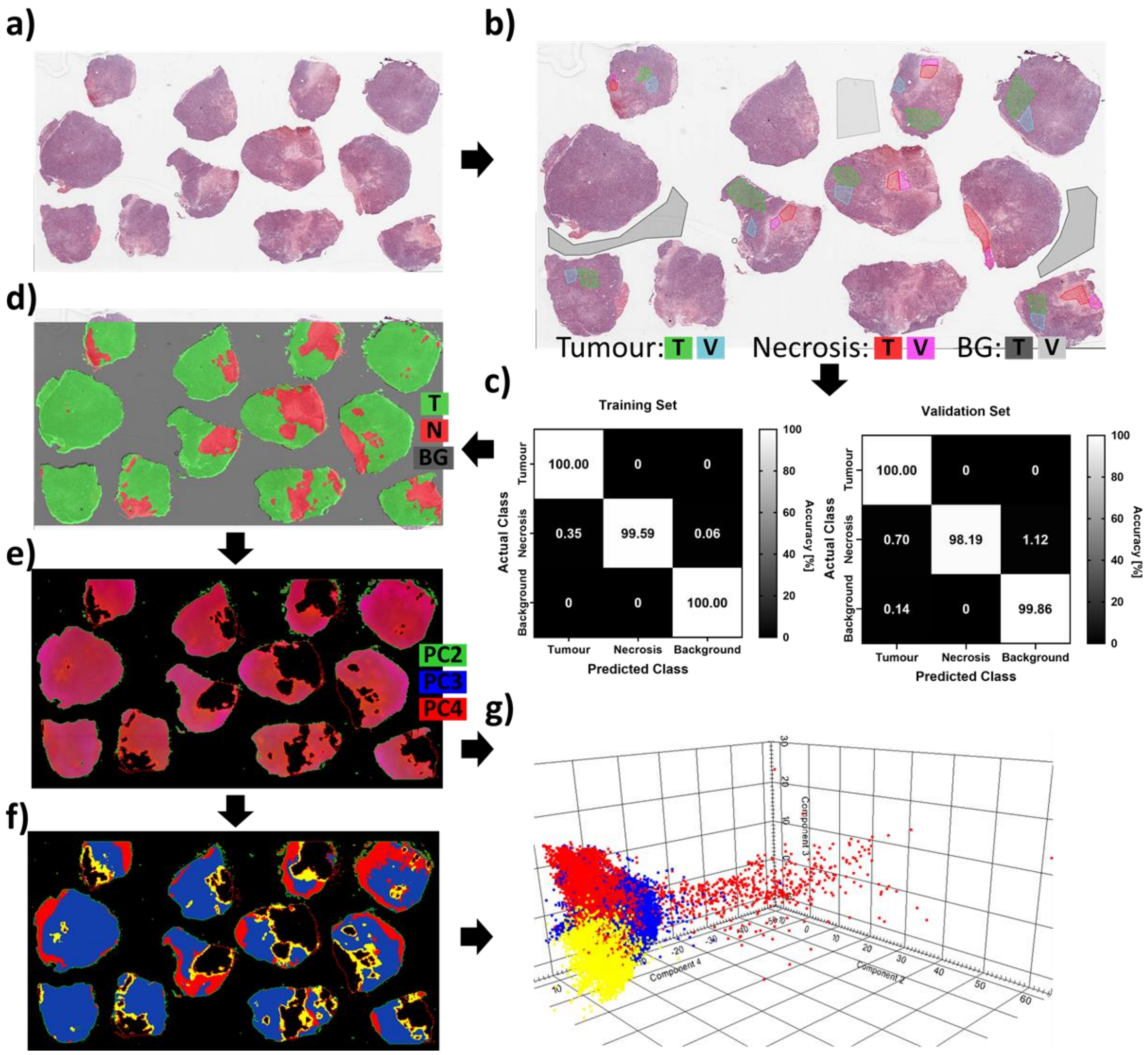
